# Cooperative super-enhancer inactivation caused by heterozygous loss of CREBBP and KMT2D skews B cell fate decisions and yields T cell-depleted lymphomas

**DOI:** 10.1101/2023.02.13.528351

**Authors:** Jie Li, Christopher R. Chin, Hsia-Yuan Ying, Cem Meydan, Matthew R. Teater, Min Xia, Pedro Farinha, Katsuyoshi Takata, Chi-Shuen Chu, Martin A. Rivas, Amy Chadburn, Christian Steidl, David W. Scott, Robert G. Roeder, Christopher E. Mason, Wendy Béguelin, Ari M. Melnick

## Abstract

Mutations affecting enhancer chromatin regulators *CREBBP* and *KMT2D* are highly co-occurrent in germinal center (GC)-derived lymphomas and other tumors, even though regulating similar pathways. Herein, we report that combined haploinsufficiency of *Crebbp* and *Kmt2d* (C+K) indeed accelerated lymphomagenesis. C+K haploinsufficiency induced GC hyperplasia by altering cell fate decisions, skewing B cells away from memory and plasma cell differentiation. C+K deficiency particularly impaired enhancer activation for immune synapse genes involved in exiting the GC reaction. This effect was especially severe at super-enhancers for immunoregulatory and differentiation genes. Mechanistically, CREBBP and KMT2D formed a complex, were highly co-localized on chromatin, and were required for each-other’s stable recruitment to enhancers. Notably, C+K lymphomas in mice and humans manifested significantly reduced CD8^+^ T-cell abundance. Hence, deficiency of C+K cooperatively induced an immune evasive phenotype due at least in part to failure to activate key immune synapse super-enhancers, associated with altered immune cell fate decisions.

**SIGNIFICANCE:** Although CREBBP and KMT2D have similar enhancer regulatory functions, they are paradoxically co-mutated in lymphomas. We show that their combined loss causes specific disruption of super-enhancers driving immune synapse genes. Importantly, this leads to reduction of CD8 cells in lymphomas, linking super-enhancer function to immune surveillance, with implications for immunotherapy resistance.

## INTRODUCTION

Epigenetic mechanisms play a central and critical role in malignant transformation and in encoding tumor phenotypes (1–3). Epigenetic gene regulation is mediated by many intersecting layers of chromatin modifying proteins, transcription factors, and other mechanisms (4). Parsing out how these various components cooperate and influence each other to yield specific tumor phenotypes, and the means for therapeutic targeting these complex mechanisms remains a critical unmet need. The fact that somatic mutations of chromatin modifiers and transcription factors are among the most common genetic events in cancer underlines the power and relevance of these mechanisms. In particular, chromatin modifier gene mutations are extremely frequent in follicular lymphoma (FL) and diffuse large B-cell lymphoma (DLBCL), the two most common non-Hodgkin lymphomas (NHLs) (5–7). Both of these tumors arise mostly from B cells transiting through the germinal center (GC) reaction during the humoral immune response (8). DLBCLs manifesting a stronger GC-like transcriptional profile (GCB-DLBCLs) harbor the highest incidence of chromatin modifier gene mutations (9).

The two most recurrently mutated genes in these tumors are *KMT2D* and *CREBBP*, which are believed to be founding events in lymphomagenesis (10, 11). KMT2D is a histone methyltransferase that selectively mediates H3K4 mono- and di-methylation (H3K4me1/me2) primarily at enhancers to prime their activation (12, 13). KMT2D is most commonly affected by generally heterozygous nonsense mutations that yield a truncated protein lacking the enzymatic SET domain. CREBBP is a histone acetyltransferase that mediates enhancer activation through acetylation of H3K27 and other residues (14–16). CREBBP loss of function in FL and DLBCL is caused by either missense mutations in the histone acetyltransferase domain that inactivate enzymatic activity, or by truncating mutations. Homozygous conditional knockout of *Kmt2d* or *Crebbp* in mouse GC B cells leads to focal reduction of H3K4me1 and H3K27ac respectively, preferentially at gene enhancers (17–22). Many of the affected genes overlap and are involved in immune signaling pathways, which play important roles in GC physiology.

GCs are dynamic microanatomical structures formed transiently upon T cell-dependent antigen challenge (23, 24). GCs feature a dark zone (DZ) consisting of densely packed and rapidly proliferating B cells called centroblasts (CBs) undergoing somatic hypermutation (SHM) of their immunoglobulin gene variable regions. Upon exhausting their resources CBs migrate to the light zone (LZ) as smaller, post-replicative centrocytes (CCs). CCs compete vigorously for T cell help from a limiting number of specialized T follicular helper cells (TFH). Depending on the strength and quality of these immune synapse interactions, CCs will adopt different cell fate decisions including: 1) recycling back to the DZ for additional rounds of division and SHM (“DZ recycling”); 2) differentiation into plasma cells (PCs); 3) transition to the various types of memory B cells (MBs); or 4) undergoing apoptosis. The proliferative and mutagenic nature of GC B cells render them highly susceptible to off-target mutation and malignant transformation. *CREBBP* and *KMT2D* loss of function was proposed to alter how GC B cells respond to T cell help by preventing enhancer activation (19–22).

It is a generally accepted principle that within a given tumor, oncogenic hits that engage the same pathways or processes would be generally mutually exclusive to each other. One example of this are the mutually exclusive mutations in TET2 and IDH enzymes in acute myeloid leukemia (25). The reason for their exclusivity is believed due to their both inducing gene promoter cytosine hypermethylation as an important part of their oncogenic functions. Likewise, mutations in *CREBBP* and *KMT2D* seem to both impair the function of highly overlapping GC exit enhancers in B cells. However, it was shown that the simultaneous presence of *CREBBP* and *KMT2D* mutations represent the most highly recurrent, paired gene co-occurrence scenario in B-cell lymphomas (26).

This apparent paradox raises many intriguing questions as to the nature of oncogenic epigenetic reprogramming in tumors, especially related to gene enhancers. For example, it was shown that combining chromatin modifier with signaling protein mutations (e.g., TET2 + FLT3) resulted in a synergistic effect on both epigenetic and transcriptional levels, that was different than those induced by either mutation alone (27). However, in this case, *CREBBP* and *KMT2D* mutations appear to have such overlapping mechanisms of action that it is hard to predict exactly what their combinatorial outcome would be. Formally demonstrating that these mutations cooperate in malignant transformation, and on what basis, is an intriguing question from both the epigenetic and biological standpoints. Herein we set out to explore these questions, which may be relevant not only to lymphomas but also to the many other tumors (cBioPortal) where these two mutations co-occur.

## RESULTS

### Combined loss of CREBBP and KMT2D accelerates onset of B-cell lymphomas with FL characteristics

The reported co-occurring pattern of *CREBBP* and *KMT2D* mutations across B-cell lymphomas prompted us to further validate those findings in publicly available patient datasets. Examining cohorts of FL and EZB/cluster3 DLBCL (the subset of DLBCL enriched for these mutations, n = 478 and 319, respectively) (5-7,9,11,19,26,28-30), we confirmed significant co-occurrence of these mutations (p values are 1.68E-6 and 1.77E-18, respectively, **Supplementary Fig. S1A**). These data led us to explore whether KMT2D and CREBBP loss of function might cooperate to drive malignant transformation of GC B cells. Given that these mutations are generally heterozygous, we focused on generating conditional double heterozygous knockout mice. For this we crossed *Crebbp* and *Kmt2d* floxed conditional knockout mice with the Cγ1-cre strain to induce their heterozygous deletion in GC B cells (31–33). These animals were further crossed to VavP-BCL2 mice (34) given that *CREBBP* and *KMT2D* mutant FLs and DLBCLs generally harbor *BCL2* translocations (35). We generated a cohort of mice with the following genotypes: *VavP-BCL2*;*Cγ1*^cre/+^;*Crebbp*^fl/+^;*Kmt2d*^fl/+^ (**BCL2+CK**), *VavP-BCL2*;*Cγ1*^cre/+^;*Crebbp*^fl/+^ (**BCL2+C**), *VavP-BCL2*;*Cγ1*^cre/+^;*Kmt2d*^fl/+^ (**BCL2+K**), *VavP-BCL2*;*Cγ1*^cre/+^ (**BCL2**), and *Cγ1*^cre/+^(**WT**) controls (**Fig.1A**). To expand the number of mice and generate age- and sex-balanced cohorts, bone marrow of donor mice with these engineered alleles was transplanted into lethally irradiated recipients (n=35 per genotype), which were subsequently immunized with the T cell-dependent antigen sheep red blood cells (SRBCs) at several intervals to induce GC formation and lymphomagenesis (**Fig. 1A**).

**Figure 1.**
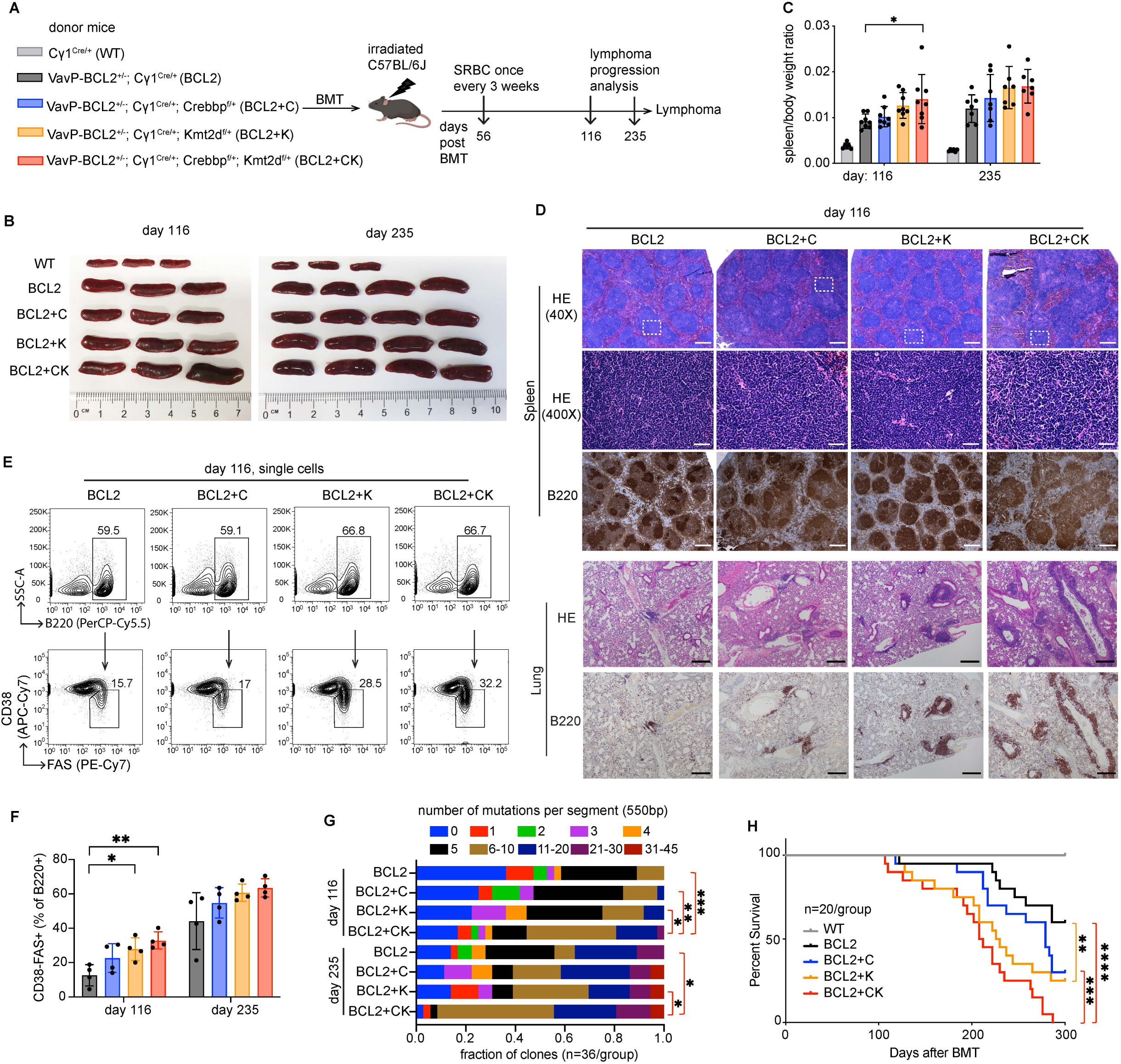
Crebbp and Kmt2d haploinsufficiency cooperatively accelerates murine lymphomagenesis. **A**, Experimental scheme for murine lymphomagenesis. **B-C**, Representative (**B**) spleen images, and (**C**) spleen/body weight ratio (mean ± SD) of mice euthanized at day 116 and 235 post-bone marrow transplantation (BMT) (n=6-8 mice per genotype). Each dot in (**C**) represents a mouse. Statistical significance was determined using ordinary one-way ANOVA followed by Tukey-Kramer’s multiple comparisons test (*p < 0.05). **D**, Representative images of hematoxylin and eosin (H&E) and B220 immunohistochemistry (IHC) staining of formalin-fixed paraffin-embedded spleen and lung sections prepared from mice euthanized at day 116 post-BMT. Bottom H&E images of spleens show the zoom-in of outlined areas in top images. The scale bars represent 200 pixels in spleen and 380 pixels in lung. **E**, Representative FACS plots show the gating strategy and relative frequency of splenic B220^+^ B cells and CD38^-^FAS^+^ GC B cells of mice at day 116 post-BMT. **F**, FACS analysis showing the relative abundance of splenic GC B cells (CD38^-^FAS^+^) as percent of B220^+^ cells (mean ± SD) at days 116 and 235 post-BMT. Each dot represents a mouse (n=4 mice per genotype). Statistical significance was determined using ordinary one-way ANOVA followed by Tukey-Kramer’s multiple comparisons test (*p < 0.05; **p < 0.01). **G**, Somatic hypermutation (SHM) profiling of the IgH-VJ558-JH4 region (550bp in length) in GC B cells sorted from spleen of mice at days 116 and 235 post-BMT. The mutation burden was determined by Sanger sequencing of PCR product covering the IgH-VJ558-JH4 region (n=36 clones per genotype). The p values were calculated by two-tailed Wilcoxon rank sum test (*p < 0.05, **p < 0.01, ***p < 0.001). **H**, Kaplan–Meier survival curve of the lymphomagenesis cohort of mice. n = 20 mice per group; the p values were calculated by two-sided log-rank test and denoted as follows: **p < 0.01, ***p < 0.001, ****p < 0.0001.

A subset of these animals was sacrificed for phenotypic characterization at day 116 post-bone marrow transplant, at which time the first overt illness was observed in mice, or at day 235, when many of the BCL2+CK mice were reaching their terminal stage (n= 6-8 per time point). At both time points, we observed progressively greater splenomegaly and spleen/body weight ratios in BCL2+C, BCL2+K, and BCL2+CK mice as compared to BCL2 mice, with significant differences observed between BCL2+CK vs BCL2 animals (**Fig. 1B-C**). Histologic analysis based on H&E and B220 staining of splenic tissue at day 116 showed well-defined although progressively hyperplastic follicular structures containing small B lymphocytes in BCL2, BCL2+C and BCL2+K mice, consistent with low-grade or incipient FL. In contrast, BCL2+CK mice manifested a mixed picture of aberrant hyperplastic follicles with markedly expanded and distorted lymphoid structures containing intermingled populations of larger B cells and tingible body macrophages, consistent with more aggressive, higher-grade FL (**Fig. 1D**). There was progressively greater abundance of lymphoma cells infiltrating solid organs in BCL2, BCL2+C, BCL2+K, and especially in BCL2+CK animals (**Fig. 1D, Supplementary Fig. S1B**). The histologic appearance of BCL2 mice at day 235 showed further enlarged follicles, composed of fairly monotonous small lymphocytes (**Supplementary Fig. S1C**). Proliferation (Ki67 staining) was relatively modest and was observed in follicular cell clusters (signal in red pulp is non-specific). A similar picture was observed in BCL2+C animals although with a few larger B cells among malignant follicles. BCL2+K animals had a small centroblastic or blastoid morphology with greater numbers of proliferating cells. BCL2+CK manifested highly enlarged and distorted lymphoid structures composed of larger, more proliferative blastoid cells as compared to the other genotypes (**Supplementary Fig. S1C**), indicating a histologically more advanced and aggressive FL.

Consistent with the FL phenotype of these tumors, flow cytometry analysis showed progressively increased proportion of GC B cells (CD38^-^FAS^+^) among total B cells (B220^+^) in the four genotypes at day 116, which had plateaued by day 235 (**Fig1. E-F, Supplementary Fig. S1D-E**). Confirming the GC origin of these tumors, there was ample evidence of somatic hypermutation (SHM) at the VDJ VJ558-JH4 locus across all genotypes (**Fig. 1G**). Of note, the SHM burden was significantly greater in BCL2+CK lymphoma cells at both time points as compared to all other genotypes (**Fig. 1G**), suggesting these cells were more exposed to DZ-like proliferative bursting and AICDA activity. Finally, an overall survival analysis showed a similar trend of progressively inferior outcome from BCL2+C, BCL2+K, to BCL2+CK compared to BCL2 (**Fig. 1H**). Taken together, these data indicate that CREBBP and KMT2D loss of function cooperates to accelerate development of FLs with more aggressive characteristics than either allele alone.

### CREBBP and KMT2D haploinsufficiency induces GC hyperplasia and superior GC B cell fitness

Given that *CREBBP* and *KMT2D* loss of function mutations are clonal in humans and cooperate in mice to induce aggressive lymphomas, we next explored whether they also interact to perturb their GC cell of origin to yield a preneoplastic phenotype. For this we generated mice with the following genotypes: *Cγ1*^cre/+^, *Cγ1*^cre/+^;*Crebbp*^fl/+^, *Cγ1*^cre/+^;*Kmt2d*^fl/+^, and *Cγ1*^cre/+^;*Crebbp*^fl/+^;*Kmt2d*^fl/+^, hereafter referred to as WT, C, K, and CK respectively. Sex and age-matched littermates were immunized with SRBCs to trigger GC formation and were euthanized 10 days later, when GCs reached their peak size (**Fig. 2A**). Flow cytometry analysis revealed that the abundance of total B cells (B220^+^) was comparable among all genotypes (**Fig. 2B**). In contrast, the proportion of GC B cells (CD38^-^FAS^+^) was progressively increased in C, K, and CK mice relative to WT, with CK mice manifesting significant and massive GC hyperplasia compared to all other genotypes (**Fig. 2C-D**). To further confirm this phenotype we performed immunofluorescence (IF) staining of spleen sections using B220-AF488 (green) and Ki67-AF594 (red), which co-label actively proliferating GC B cells (**Fig. 2E**). Quantification of these images showed significant increase in the size of GCs in CK mice, with C and K manifesting intermediate sizes (**Fig. 2F**). The increased GC B cell numbers observed by flow cytometry was due to the larger size of GCs, since there was no change in the overall abundance of these structures (**Fig. 2G**). Among GC B cells, the ratio of centroblasts (CXCR4^hi^CD86^lo^) to centrocytes (CXCR4^lo^CD86^hi^) was significantly skewed towards centroblasts in C and CK (**Fig. 2C, 2H**).

**Figure 2.**
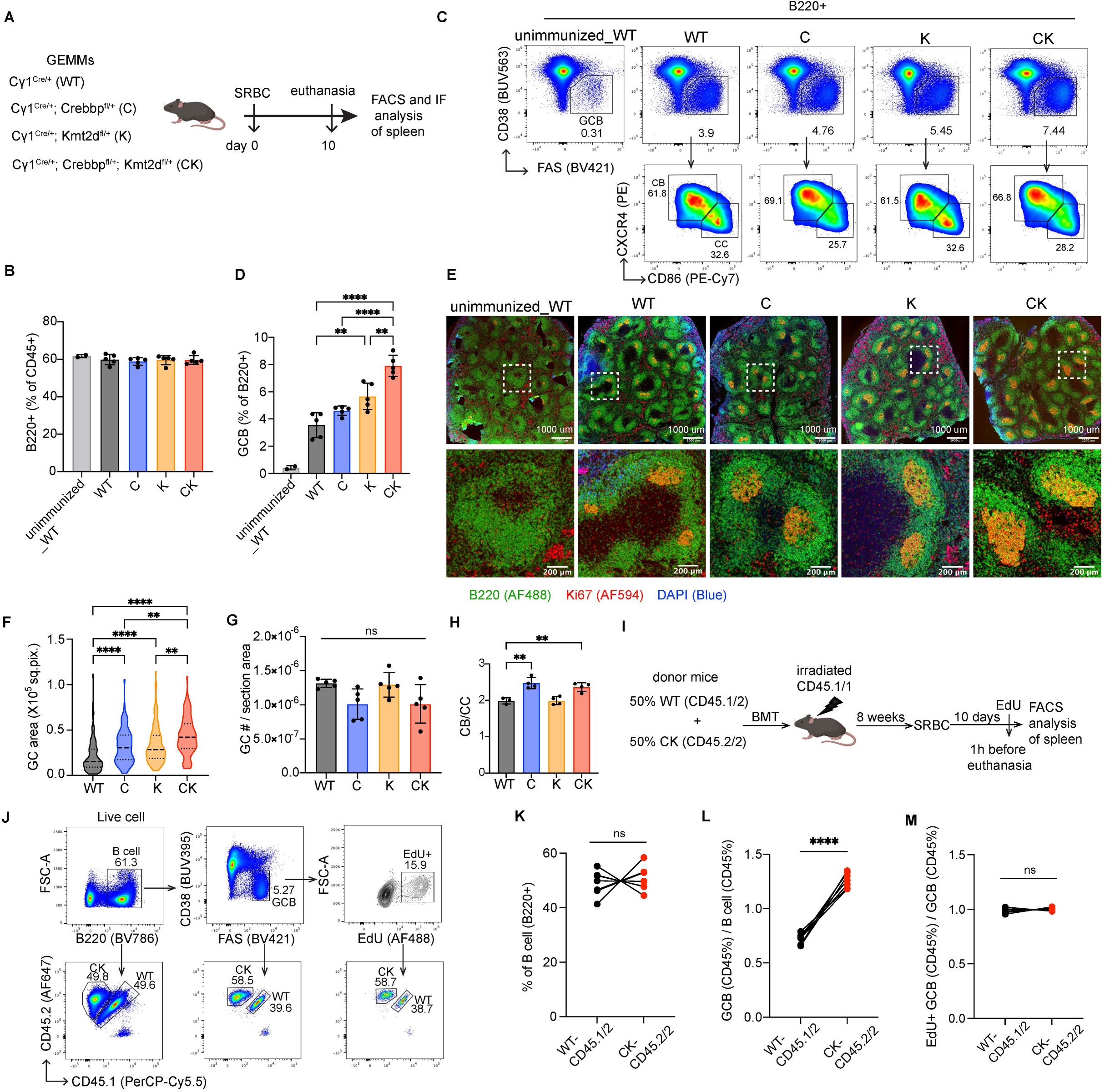
Crebbp and Kmt2d haploinsufficiency cooperatively induces hyperplastic GCs with superior fitness. **A**, Experimental scheme for GC characterization (results shown in **B-H**). IF: immunofluorescence. **B,D,H,** FACS analysis showing the relative abundance (mean ± SD) of splenic (**B**) total B cells (%CD45^+^ with B220^+^), (**D**) GC B cells (%B220^+^ with CD38^-^FAS^+^), and (**H**) CB (%GC B with CXCR4^hi^CD86^lo^) divided by CC (%GC B with CXCR4^lo^CD86^hi^). Each dot represents a mouse (n=2-5 per genotype). Statistical significance was determined using ordinary one-way ANOVA followed by Tukey-Kramer’s post-test (**p < 0.01, ****p < 0.0001). **C**, Representative FACS plots show the gating strategy and relative frequency of splenic CD38^-^FAS^+^ GC B cells, CXCR4^hi^CD86^lo^ CB and CXCR4^lo^CD86^hi^ CC. **E**, B220 (AF488, green), Ki67 (AF594, red), and DAPI (blue) IF staining images of spleen sections. Bottom images show the zoom-in of outlined areas in top images. Scale, 1000 um (top), 200 um (bottom). **F-G**, Quantification of (**F**) splenic GC area (left to right: n=119, 91, 122, 95 GCs) and (**G**) relative splenic GC number normalized to spleen section area (each dot represents a mouse, mean ± SD) based on IF images as shown in (**E**). Statistical significance was determined using Kruskal-Wallis test followed by Dunn’s multiple comparisons test (**F**) or ordinary one-way ANOVA followed by Tukey-Kramer’s post-test (**G**) (ns p > 0.05, **p < 0.01, ****p < 0.0001). **I**, Experimental design for fitness study (results shown in **J-M**). **J**, Representative FACS plots show gating strategy and relative frequency of the indicated splenic cell types (left to right: total B, GC B, EdU+ GC B cells). WT and CK-derived cells were separated as CD45.1/2 and CD45.2/2, respectively. **K**, FACS data showing the proportion of WT (CD45.1/2, black dots) and CK (CD45.2/2, red dots)-derived splenic total B cells (B220^+^) in recipients. Each pair of connected dots represents a mouse (n=7). P value was calculated by two-tailed paired t test (ns p > 0.05). **L-M**, FACS data showing the ratio of WT (CD45.1/2, black dots) or CK (CD45.2/2, red dots)-derived GC B cell percentage to their respective parental total B cell percentage (**L**) or EdU+ GC B cell percentage to their respective parental total GC B cell percentage (**M**). Each pair of connected dots represents a mouse (n=7). P value was calculated by two-tailed paired t test (ns p > 0.05, ****p < 0.0001).

Given this hyperplastic phenotype, we next generated bone marrow chimeric mice to evaluate whether CK GC B cells exhibit a competitive advantage over WT controls when placed within the same microenvironment. Bone marrow cells carrying distinct congenic markers WT: CD45.1/2 and CK: CD45.2/2, were mixed at equal ratio before transplanting into lethally irradiated recipients (CD45.1/1). Fully engrafted mice were immunized with SRBCs and euthanized 10 days later, 1 hr after injection of EdU (**Fig. 2I**). As expected, flow cytometry analysis of splenic cells revealed that WT and CK-derived cells were equally represented in the total B cell population (**Fig. 2J-K**). By contrast, normalizing the percentage of WT or CK in GC B cells to their respective percentage in total B cells we observed a significantly increased proportion of CK-derived GC B cells vs WT controls (**Fig. 2J, 2L**). This increased competitiveness was not due to changes in proliferation rate as shown by EdU incorporation (**Fig. 2J, 2M**). Collectively, these data suggest that simultaneous monoallelic deficiency of CREBBP and KMT2D confers GC B cell fitness advantage without changes in cell proliferation rate.

### CREBBP and KMT2D haploinsufficiency induces cooperative disruption of GC transcriptional programming

To dissect the molecular mechanisms underlying the combinatorial perturbation of the GC reaction by CREBBP and KMT2D heterozygosity, we performed RNA-seq profiling of CB and CC sorted from SRBC-immunized mice from each genotype (**Fig. 3A**). Principal component analysis (PCA) showed the expected clear separation between CB and CC along the PC1 axis (**Fig. 3B**). Among CBs, only CK (but not C or K) cells manifested a clear difference compared to WT along the PC2 axis. CK CC transcriptional profiles were also markedly distinct from WT, although in this case the K cells reflected an intermediate degree of perturbation. Similarly, hierarchical clustering revealed clear segregation of CK CBs and CCs from the two clusters formed by WT/C/K-CBs and WT/C/K-CCs (**Fig. 3C**). To better understand these observations we performed a supervised analysis comparing each mutant genotype to WT, to define differentially expressed genes among CBs and CCs. This analysis revealed significantly greater perturbation among CK cells, with transcriptional changes skewed towards repression (295 genes up and 1147 genes down in CB, 367 genes up and 1417 genes down in CC, q<0.01 and |FC|>1.5). Many of these genes were also perturbed by C or K, albeit to a generally lower degree (**Fig. 3D-E**, **Supplementary Fig. S2A-B**, **and Supplementary Table 1-2**). A hypergeometric pathway analysis enriched for gene signatures similarly altered in CK CBs and CCs (**Fig. 3F, Supplementary Table 3**). In particular, gene sets associated with GC exit signaling, DNA damage repair, and tumor suppressors were expressed at lower levels in CK relative to WT (**Fig. 3F**). Conversely, those related to biosynthetic metabolism were upregulated in CK compared to WT (**Fig. 3F**). Altogether, these data support the notion that CREBBP and KMT2D dual haploinsufficiency results in cooperative repression of GC exit/immune synapse responsive genes and DNA repair, which is consistent with skewing towards the CB transcriptional state. Simultaneous induction of biosynthetic programs normally restricted to CCs undergoing T cell help may further explain the competitive advantage of these cells (36).

**Figure 3.**
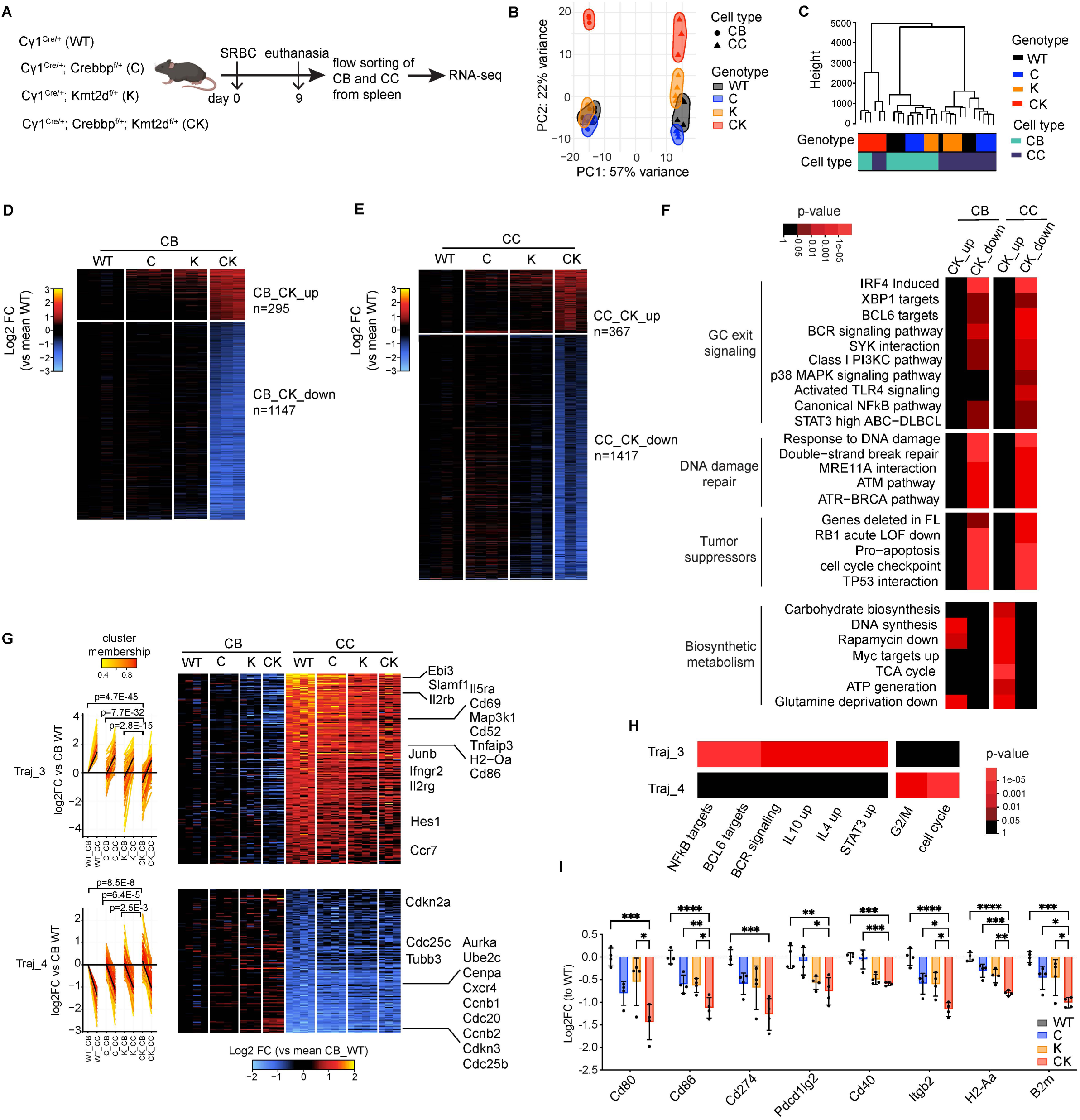
RNA-seq reveals cooperative perturbation of intra-GC transcriptional transitions by Crebbp and Kmt2d haploinsufficiency. **A**, Experimental scheme for RNA-seq profiling of CB and CC (results shown in **B-H**). **B**, Principal component analysis (PCA) of RNA-seq datasets using all genes normalized as VST (variance stabilizing transformation) (n=3-4 mice per genotype). **C**, Dendrogram showing the hierarchical clustering result of RNA-seq datasets using the top 10% most variable genes normalized as VST, the Manhattan distance and ward.D2 linkage. **D-E**, Heatmap showing the relative expression levels of the union of differentially expressed genes (DEGs) as log2-transformed ratio of every sample to the mean of wild-type gene expression for each genotype in (**D**) CB and (**E**) CC. Each column represents one mouse dataset. Union DEGs include all DEGs defined in at least one pair-wise comparison using WT as control with a significance cut-off of FDR<0.01, |log2FC|>0.58. **F**, Pathway enrichment analysis for CK vs WT DEGs using Parametric Analysis of Gene Set Enrichment (PAGE). P values were calculated by hypergeometric mean test. **G**, Fuzzy c-means clustering of RNA-seq datasets identified 8 clusters (named as Traj_1 to Traj_8) with distinct trajectory pattern, Traj_3 and Traj_4 are shown: line plot (left) and heatmap (right) of standardized log2 fold-change relative to mean of WT CB. Black lines in the line plot are cluster centroid; each gene is colored by the degree of cluster membership. A linear regression model was fit for log2 fold-change expression compared to WT as a function of cell type (CB or CC), genotype (C, K, or CK), and interaction of cell type and genotype. The corresponding p values for the coefficients are shown. **H**, Pathway enrichment analysis using PAGE for Traj_3 and Traj_4 genes defined in panel **G**. P values were calculated by hypergeometric mean test. **I**, RT-qPCR of indicated genes in sorted GC B cells for each genotype (n = 4 mice per genotype). Each dot represents a mouse. qPCR signal for each gene was normalized to those of GAPDH and then mean WT and presented as log2 fold-change ± SEM. Statistical significance was determined using ordinary one-way ANOVA followed by Tukey-Kramer’s multiple comparisons test (*p < 0.05, **p < 0.01, ***p < 0.001, ****p < 0.0001).

To gain further insight into how CB-to-CC transcriptional reprogramming is altered in CK-deficient GCs, we conducted trajectory analysis of the gene expression change from CB to CC for each genotype, and then grouped these differentially expressed genes into eight distinct clusters (named as Traj_1 to Traj_8) by Fuzzy c-means clustering (**Supplementary Fig. S2C**). Trajectories 3 (Traj_3) and 4 (Traj_4) were most informative. Traj_3 genes are normally upregulated when CBs transition to CCs, and displayed progressively lower basal expression in CBs and impaired upregulation in CCs across C, K and CK (**Fig. 3G**). Reciprocally, Traj_4 genes that are repressed in WT CCs vs CB, were expressed at higher baseline levels and were progressively less repressed across C, K and CK genotypes (**Fig. 3G**). Of particular interest, Traj_3 included CC-signature immune synapse responsive genes such as *Cd86* and *H2-Oa*, whereas Traj_4 contained genes *Cxcr4* and *Ccnb1*, involved in DZ GC B cells and their classical G2/M cell cycle progression phenotype (**Fig. 3G**, **Supplementary Table 1-2**). Consistent with these findings, Traj_3 was significantly enriched with NFkB and BCL6 target genes, as well as genes linked to BCR, IL10, IL4 and STAT3 signaling. These data are consistent with the known role of CREBBP and KMT2D in antagonizing BCL6-mediated gene repression (19) (**Fig. 3H, Supplementary Table 3**). Traj_4 enriched genes included cell cycle and G2/M gene sets, consistent again with the strengthened DZ signature. We validated these results in independent experiments using RT-qPCR to measure abundance of canonical CC-induced immune synapse genes such as *Cd86* and *H2-Aa* (**Fig. 3I**). Taken together, these results demonstrate that CREBBP and KMT2D haploinsufficiency cooperatively attenuates induction of genes essential for CB-to-CC cell state transition.

### CREBBP and KMT2D haploinsufficiency skews GC B cell fate decisions towards DZ vs GC exit

The transcriptional programming effects noted above suggested that CK deficiency might alter the flux of CCs towards their various downstream cell fates. Given that GC B cells consist of a heterogenous continuum of subpopulations beyond the resolution of standard flow cytometry methods, we used single cell RNA-seq (scRNA-seq) to determine the impact of C, K and CK haploinsufficiency on the abundance of GC B subpopulations. To accomplish this, we sorted splenic B220^+^IgD^-^ B cells from each genotype at day 10 post-SRBC immunization, followed by scRNA-seq. Uniform manifold approximation and projection (UMAP) was applied for dimensionality reduction, then cell subpopulations were identified using Graph-based clustering and K nearest neighbor (**Fig. 4A**). No sample specific batch effects were observed (**Supplementary Fig. S3A**). GC B cell clusters were subsequently annotated as early_centroblast (early_CB), centroblast, transitioning_CB_CC (trans_CB_CC), centrocyte, recycling, and pre-memory by label transfer from previously published data sets (37). The accuracy of these assignments was further validated using cell type-specific gene signatures and marker genes (**Fig. 4A-B, Supplementary Fig. S3B**). Differential cell abundance analysis revealed that the relative proportion of early_CB and CB were progressively increased, whereas that of centrocyte and prememory B cell were progressively decreased in C, K, and CK as compared to WT (**Fig. 4C**).

**Figure 4.**
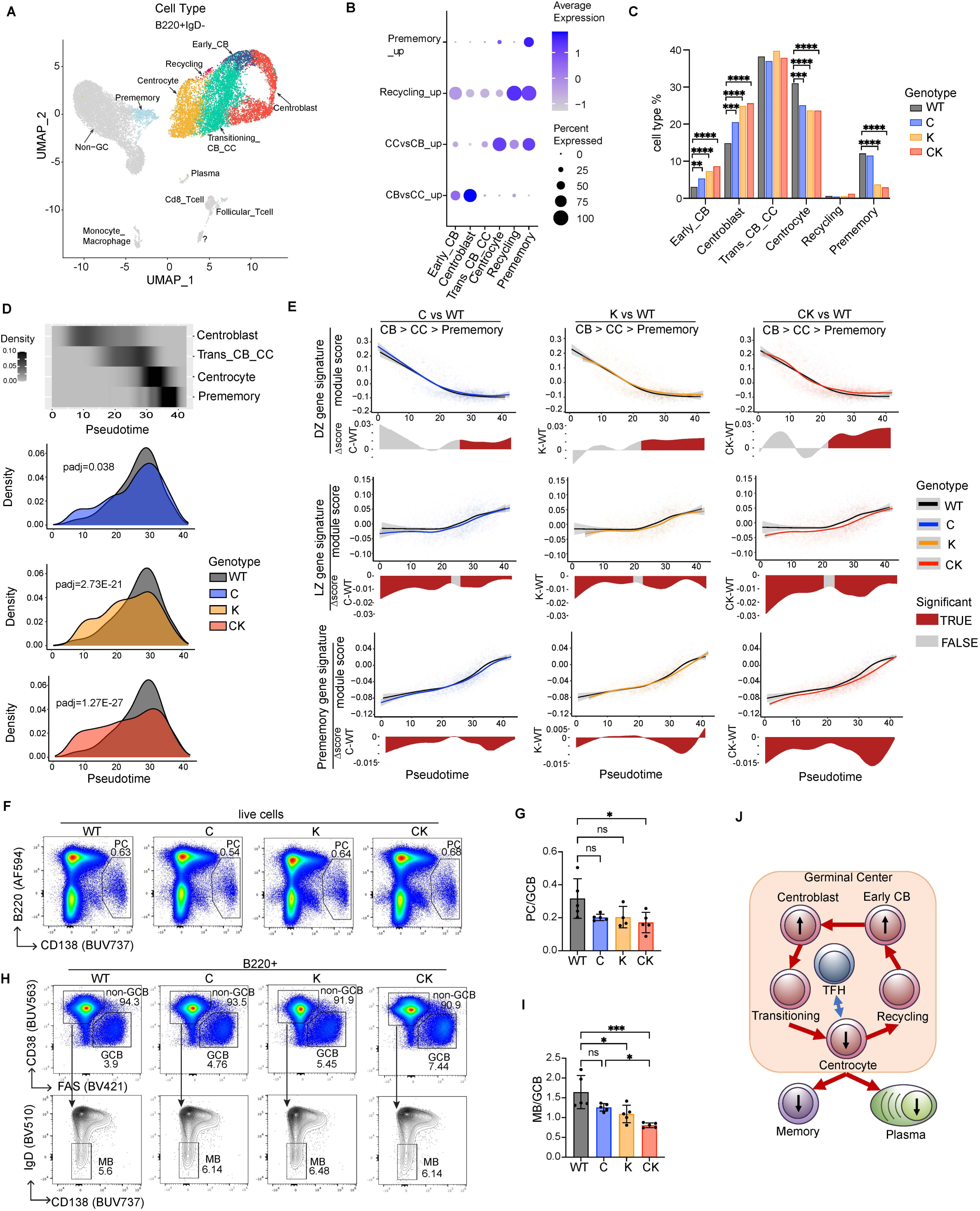
Crebbp and Kmt2d haploinsufficiency cooperatively skews the GC B cell fate toward CB over exit differentiation into MB and PC. **A-E**, Splenic B220^+^IgD^-^ cells sorted from mice immunized with SRBC for 10 days were subject to single-cell RNA-seq for each genotype (WT, C, K, and CK). **A**, UMAP representation of the single-cell RNA-seq dataset, with the highlighted GC B cell subtypes annotated by label transfer from previously published datasets (37). Grayed cell clusters are either non-GC B cells or other contaminating cell types. **B**, Dot plot in which the color intensity and shape size of each dot corresponds to the average gene expression and percent of cells expressing the signature, respectively. **C**, Bar plot depicting the relative proportion of the indicated GC B cell subtypes in each genotype. Statistical significance was determined using Fisher’s exact test (**p < 0.01, ***p < 0.001, ****p < 0.0001). **D**, Cell density plot showing the frequency of the indicated GC B cell subtypes along the Slingshot pseudotime axis anchored at CB. The pair-wise differential cell density distribution was assessed by two-sided Wilcoxon test. **E**, Top, module score of the DZ, LZ, or prememory gene signatures was plotted for each cell on the y axis, with a best fit spline curve representing the average score. Bottom, cells were binned into ten bins with the same number of cells per bin, and differential expression shown as a delta spline plot across pseudotime, colored based on statistical significance determined by two-sided Wilcoxon test for each bin. **F, H**, Representative FACS plots show the gating strategy and relative frequency of splenic (**F**) plasma cells (PC, B220^lo^CD138^+^) and (**H**) memory B cells (MB, CD38^+^FAS^-^ IgD^-^CD138^-^) at day 10 post-SRBC. **G, I**, FACS analysis showing (**G**) the PC/GC B percentage ratio, both normalized to total live cells, and (**I**) MB/GC B percentage ratio, both normalized to total B cells at day 10 post-SRBC (mean ± SD). Each dot represents a mouse. Statistical significance was determined using ordinary one-way ANOVA followed by Tukey-Kramer’s post-test (ns p > 0.05, *p < 0.05, ***p < 0.001). **J**, Graphical representation depicting the cell state transitions within GC. Upward and downward black arrows indicate increase and decrease, respectively, of the associated cell type abundance in CK-deficient GC compared to WT.

To further assess these changes, we used slingshot (38) to generate a pseudotime trajectory starting in CB, passing through the Trans_CB_CC and CC, and ending at prememory cells (**Fig. 4D**). Plotting cell density for each genotype along this pseudotime axis showed progressively more significant shifts in distribution towards the CB side in C, K and CK (**Fig. 4D**). We next projected DZ, LZ, and prememory gene signature scores (module scores) along the pseudotime axis and assessed whether their pseudo-temporal expression patterns are altered. Notably, DZ signature genes were aberrantly maintained at higher levels in CC and prememory cells from C, K and to a greater degree CK compared to WT (**Fig. 4E**). Conversely, LZ and prememory signature genes were expressed at lower levels in C, K and to a greater degree CK, starting at CB and persisting throughout the entire differentiation trajectory, implying that C, K, and especially CK-deficient GC B cells fail to properly engage GC exit genes and may have impaired GC exit capabilities (**Fig. 4E**). Indeed, measuring abundance of memory and plasma cells relative to GC B cells by flow cytometry revealed significant reduction in these populations that was most severe in CK-deficient mice (**Fig. 4F-I**).

The relative reduction in plasma cells led us to explore whether CK deficiency has any effect on antibody affinity maturation and long-lived plasma cells. For this we first examined serum antibody titers in 4-hydroxy-3-nitrophenylacetyl (NP)-ovalbumin (OVA) immunized mice at four different time points, including 70 days post-immunization when antibody secretion comes from terminally differentiated long-lived plasma cells in bone marrow (**Supplementary Fig. S3C**). ELISA assays showed significant reduction of both low (NP28) and high (NP8) affinity NP specific IgG1 at 70 days in CK mice, although there was no change in the ratios of high to low affinity (**Supplementary Fig. S3D-F**). Accordingly, analyzing bone marrow from these animals at day 85 post-immunization by ELISPOT showed a trend towards reduced NP8 (high affinity) IgG1 secreting long-lived plasma cells in CK mice (**Supplementary Fig. S3G-H**). These data suggest that although affinity maturation is unperturbed, the abundance of GC derived antibodies is reduced perhaps due to the observed GC exit impairment. Finally, we noted that altered abundance of CBs and CCs was not exactly the same by flow cytometry (**Fig. 2H**) vs scRNA-seq, perhaps because expression of key markers such as *Cd86* and others are perturbed, underlining the importance of using orthogonal methods to measure GC cell populations. Collectively, these data suggest that CREBBP and KMT2D haploinsufficiency cooperatively impairs GC B cell fate decisions, altering their phenotypic transitions to favor the DZ mutagenic state, while impairing commitment to memory or plasma cell fates (**Fig. 4J**). This effect is consistent with the heavier somatic hypermutation burden observed in CK lymphomas vs controls (**Fig. 1G**).

### CREBBP and KMT2D haploinsufficiency cooperatively blocks dynamic activation of enhancers required for intra-GC cell state transition

Considering the known functions of CREBBP and KMT2D in transcriptional activation through epigenetic modulation of chromatin, we next asked whether the above-observed CK deficiency-induced cooperative perturbation of gene expression in GC B cells is reflected at the chromatin accessibility level. To address this question, we performed ATAC-seq profiling of GC B cells purified from SRBC-immunized mice. Both PCA and unsupervised hierarchical clustering revealed clear segregation of the four genotypes (**Fig. 5A-B**). In these analyses C was clearly different from WT, whereas K and CK were highly distinct to WT and C, but more related to each other. We next identified differentially accessible peaks for each mutant genotype relative to WT. K-means clustering grouped these peaks into three distinct clusters: K_CK_Loss regions (n=218) which manifested similar reduction in K and CK vs WT and C, while CK_Gain (n=1153) and CK_Loss (n=828) showed progressive gain or loss of signal respectively from C, K to CK relative to WT (**Fig. 5C-D, Supplementary Table 4**). Genomic feature annotation showed that CK_Gain peaks were significantly enriched at promoters and significantly depleted at enhancers. In contrast, K_CK_Loss peaks were depleted from promoters but significantly enriched at enhancers, whereas the CK_Loss peaks were enriched both at enhancers and super-enhancers (**Fig. 5E**). Accordingly, splitting all peaks into bins based on their accessibility fold change showed progressively greater representation of enhancers at sites with greater loss of ATAC-seq reads (**Fig. 5F**).

**Figure 5.**
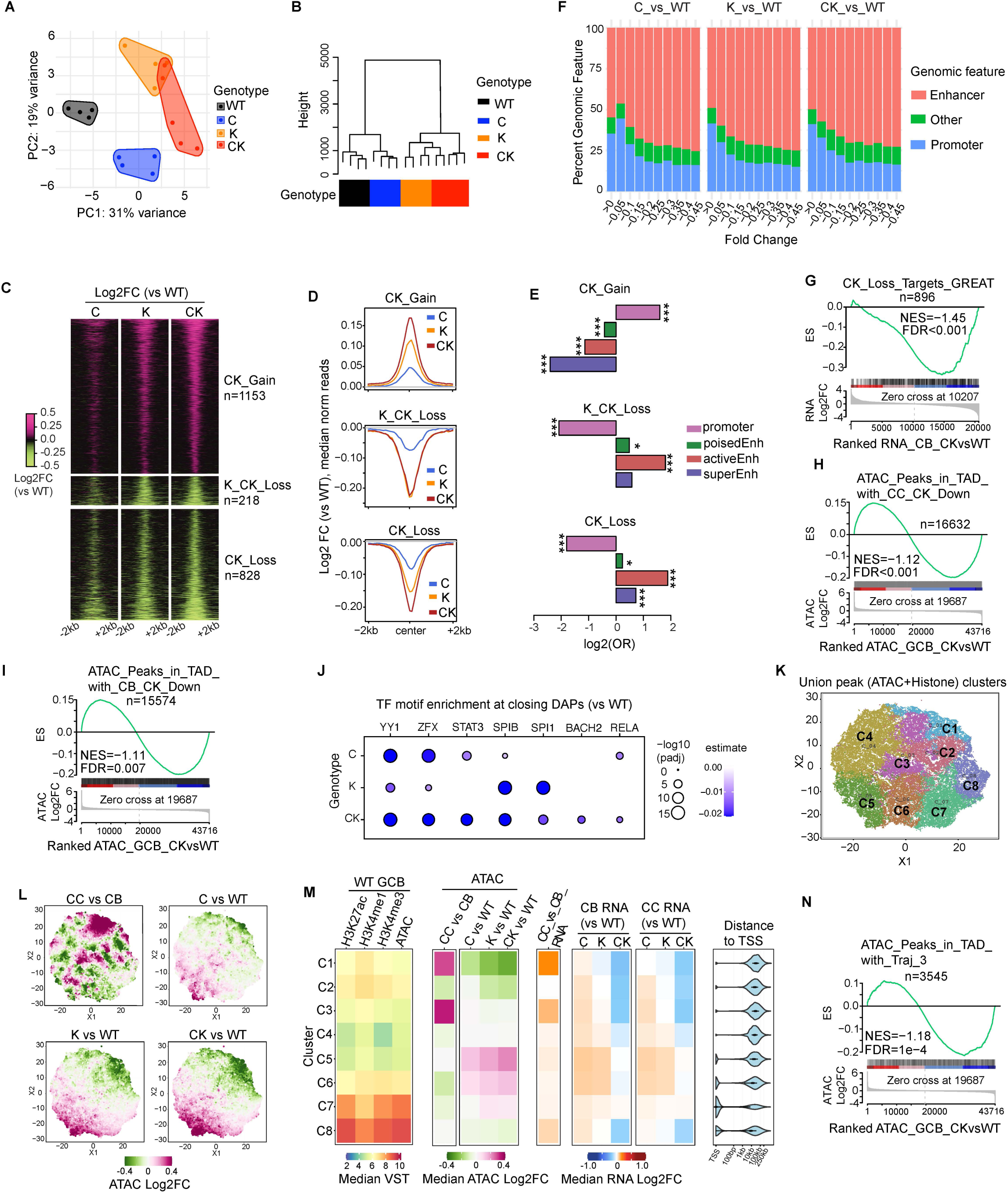
Crebbp and Kmt2d haploinsufficiency cooperatively disrupts dynamic enhancer accessibility remodeling required for intra-GC cell state transitions. **A**, PCA analysis of mouse GC B cell ATAC-seq datasets using all peaks normalized by VST (n=4-5 mice per genotype). **B**, Dendrogram showing the hierarchical clustering by ward.D2 based on Manhattan distance of mouse GC B ATAC-seq datasets using the top 10% most variable peaks normalized by VST. **C**, K-means clustered heatmap showing the relative ATAC-seq read density of the union of differentially accessible peaks (DAPs). Union DAPs include all DAPs defined in at least one pair-wise comparison using WT as control with an adjusted significance cut-off of 0.01. Three clusters (named as CK_Gain, K_CK_Loss, and CK_Loss) with distinct patterns were identified. **D**, Median ATAC-seq read intensity around peak center (+/-2kb) for each cluster as shown in panel **C**. **E**, Odds ratio (OR)-based association analysis between each DAP cluster and the indicated genomic features. Promoters (n=25,991) were annotated as TSS+/-5kb, poised (n=102,493, H3K4me1 peaks that don’t overlap with H3K27ac, promoter, and super-enhancer) and active (n=20,829, H3K27ac peaks that overlap with H3K4me1, but not promoter and super-enhancer) enhancers were defined according to mouse GC B cell H3K4me1 and H3K27ac Mint-ChIP signal, GC B-specific super-enhancers (n=755) were called by ROSE (72) using the H3K27ac Mint-ChIP signal. Fisher’s exact test was applied to determine the statistical significance (*p < 0.05, ***p < 0.001). **F**, Peaks in C, K, and CK compared to WT were split into 10 different bins based on the direction and degree of peak intensity change. Stacked bar plots showing the genomic feature distribution of DAPs in each bin. Genomic features were defined based on distance to TSS (+/-3kb for promoter), inclusion in the 5’ or 3’ UTR (other). Introns, exons and all other peaks were considered as putative enhancers. **G**, GSEA analysis using CK_Loss ATAC-seq peaks-targeting genes (annotated by GREAT) as the gene set against a ranked gene list based on CK_CB vs WT_CB RNA-seq. NES, normalized enrichment score. The p value was calculated by an empirical phenotype-based permutation test. The FDR is adjusted for gene set size and multiple hypotheses testing. **H-I**, GSEA analysis using ATAC-seq peaks in topologically associating domains (TADs) containing CK_vs_WT downregulated genes in (**H**) CC or (**I**) CB as peak set against ranked ATAC-seq peak list based on CK vs WT GC B cells. The p value was calculated by an empirical phenotype-based permutation test. The FDR is adjusted for gene set size and multiple hypotheses testing. **J**, Dot plot showing that binding motifs of the indicated TFs are enriched at closing ATAC-seq peaks in C, K or CK relative to WT. A multivariate TF regulatory potential model was employed to identify the enriched TFs. Dots are colored by accessibility remodeling score. P values were calculated by two-sided t test and corrected by Benjamini-Hochberg for multiple hypotheses testing. **K**, t-SNE dimensionality reduction and K-means clustering of union peaks, generated by merging WT GC B H3K4me3, H3K4me1, and K3K27ac Mint-ChIP datasets along with ATAC-seq datasets for each genotype (WT, C, K, and CK), produced eight distinct clusters (named as C1-C8). **L**, t-SNE plot view of the indicated relative ATAC-seq read density. The projected values are log2FC signals between the indicated populations for each peak. **M**, Heatmaps showing median VST normalized read density of the indicated histone marks or ATAC-seq for each cluster (left), or relative (median log2FC) ATAC-seq (middle) and RNA-seq (right) signal between indicated comparisons for each cluster. Distance to TSS plot shows the distance of union peaks to their closest TSSs. **N**, GSEA analysis using ATAC-seq peaks in TADs containing Trajectory_3 genes (Fig. 3G) as peak set against ranked ATAC-seq peak list based on CK vs WT GC B cells. The p value was calculated by an empirical phenotype-based permutation test. The FDR is adjusted for gene set size and multiple hypotheses testing.

Integrating our RNA-seq and ATAC-seq using GSEA indicated that CK_Loss target genes (n=896, annotated by closest TSS [transcription start site] using GREAT) were expressed at significantly lower levels in CK CB and CC compared to WT controls (**Fig. 5G, Supplementary Fig. S4A**). Considering the general co-localization of enhancers with their target genes within the same Topologically Associating Domains (TADs) (39), we further evaluated whether GC B cell TADs (40) with CK-down genes are enriched with closing ATAC-seq peaks. Indeed we observed significant reduction of ATAC-seq signal at peaks within TADs containing CK-repressed genes in CK vs WT GC B cells (**Fig. 5H-I**). Examining which transcription factor (TF) motifs are enriched at CK suppressed peaks yielded enrichment for immune synapse responsive TFs SPIB, SPI1, as well as STAT3 (41, 42), which was selective for CK but not K (**Fig. 5J**). There was also enrichment for YY1 and ZFX, both of which are implicated in GC formation but less well studied (43, 44). Expression of all these TFs were uniform across the four genotypes (**Supplementary Fig. S4B**), ruling out a change in expression of these factors driving these results.

Finally, to gain more insight into the relationship of CK-regulated enhancers with CB-to-CC cell state transitions, we performed t-SNE dimensionality reduction and K-means clustering analysis of all union peaks, generated by merging our ATAC-seq peaks with mouse GC B cell H3K27ac (40), H3K4me1, H3K4me3 Mint-ChIP peaks and CB/CC ATAC-seq peaks. These union peaks segregated into eight distinct clusters based on their read density pattern across different genotypes, histone marks, and chromatin accessibility (**Fig. 5K**). Projection of fold change in accessibility onto a tSNE plot allowed visualization of peaks with increased accessibility in CCs compared to CBs. Many of these overlapped with peaks that are progressively lost in C, K and CK relative to WT, and these peaks mainly localize to cluster 1 (**Fig. 5K-L**).

Further analysis of the peak clusters showed that Cluster 1 loci uniquely gained chromatin accessibility in CCs vs CBs, and at the same time became progressively less accessible from C, K to CK relative to WT (**Fig. 5M**, ATAC column). The respective genes (based on GREAT) were likewise upregulated in CCs vs CBs and progressively repressed in C, K and CK. In normal GC B cells these sites were generally marked by H3K27ac, H3K4me1 and H3K4me3. Cluster 1 loci were quite distal from TSS, suggesting that they correspond largely to enhancers (**Fig. 5M**, distance to TSS column). In agreement, most ATAC-seq peaks in TADs containing Traj_3 genes, which are normally upregulated during CB-to-CC transition, significantly lose accessibility in CK relative to WT (**Fig. 5N**). There were partially overlapping features in cluster 2 and 3, which may be due to the complexity of assigning regulatory elements to individual TSS, as well as the presence of certain bona fide CK responsive elements in these clusters. Overall, CREBBP and KMT2D haploinsufficiency was observed to cooperatively and selectively restrict dynamic activation of enhancers governing CB-to-CC cell state transition. Such selectivity may be conferred by specific sensitivity of certain GC cell fate-determining enhancer-regulating TFs to the dosage reduction of CREBBP and KMT2D.

### CREBBP and KMT2D form a complex, with interdependent chromatin modifying functions at immune synapse genes

To further explore the nature of CREBBP and KMT2D cooperation in human lymphoma cells and in a more tractable system we generated several isogenic clones of the GCB-DLBCL cell line OCI-Ly7 as follows: *CREBBP*^R1446C^ (C, a HAT inactivating mutation often found in human lymphomas), *CREBBP*-KO (CKO), *KMT2D*-KO (K), *CREBBP*^R1446C^+*KMT2D*-KO (CK), as well as CRISPR non-edited control clones (WT) (**Supplementary Fig. S5A-E**). RNA-seq profiling of WT, C, K and CK clones revealed that the CK combination perturbs transcriptional programming to a greater extent (PC1) and in a different manner (PC2) compared to C or K alone (**Fig. 6A**), akin to what was observed in mice. Genes downregulated in CK cells (n=1672) were significantly repressed in primary CK-mutant human GCB-DLBCLs vs GCB-DLBCLs lacking C, K or EZH2 mutations (referred to as epigenetic WT) (**Fig. 6B**). Reciprocally, genes upregulated in CK cells (n=1282) were also expressed at higher levels in primary human CK GCB-DLBCL (**Supplementary Fig. S5F**). The same was observed when performing GSEA on the set of genes either down or up regulated in primary human CK GCB-DLBCL vs the ranked isogenic cell gene expression profiles (**Supplementary Fig. S5G-H**). Hence our isogenic cells reflected the perturbations observed in primary DLBCL cases.

**Figure 6.**
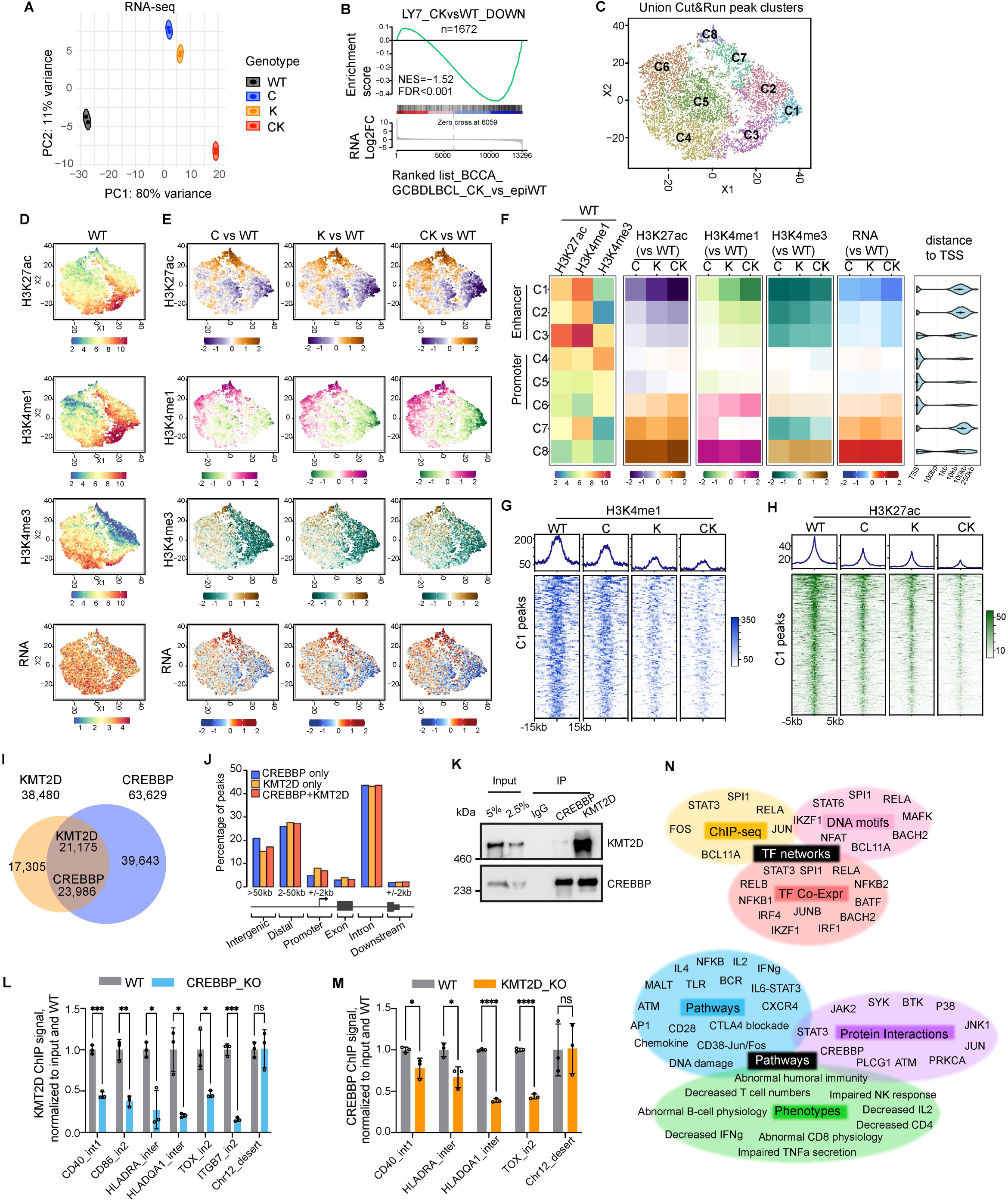
CREBBP and KMT2D form a complex, in which they reciprocally regulate each other’s chromatin binding and histone modifying activity. **A**, PCA analysis of RNA-seq datasets done in isogenic human OCI-Ly7 GCB-DLBCL cell lines (n=2 per genotype) using all genes normalized by VST. **B**, GSEA plot using CK vs WT downregulated genes in OCI-Ly7 as gene set against ranked gene list based on CK vs epigenetic WT (epiWT, no mutations in *CREBBP*, *KMT2D*, and *EZH2*) RNA-seq datasets of human BCCA cohort GCB-DLBCL patients. NES, normalized enrichment score. The p value was calculated by an empirical phenotype-based permutation test. The FDR is adjusted for gene set size and multiple hypotheses testing. **C**, t-SNE dimensionality reduction and K-means clustering of union peaks, generated by taking the union of H3K4me3, H3K4me1, and K3K27ac CUT&RUN signals from all isogenic OCI-Ly7 cell lines (WT, C, K, and CK), produced eight distinct clusters (named as C1-C8). **D**, t-SNE plots showing the basal level and distribution pattern of the indicated histone marks or RNA in WT OCI-Ly7 cells. The projected values are VST normalized counts. **E**, Read density changes for the indicated histone marks and RNA were projected onto t-SNE plots. The projected values are log2FC signal between the indicated genotypes. **F**, Heatmaps showing median VST normalized read density (left) or read density change (median log2FC) in C/K/CK relative to WT (middle and right) of the indicated histone marks or RNA-seq for each cluster defined in panel **C**. Distance to TSS plot shows the distance of union peaks to their closest TSSs. **G-H**, Average signal profiles (top) and heatmaps (bottom) displaying (**G**) H3K4me1 and (**H**) H3K27ac CUT&RUN signals around peak summit (+/-15kb) or peak center (+/-5kb), respectively at C1 peak regions. **I**, Venn diagram displaying overlap between CREBBP and KMT2D ChIP-seq peaks in OCI-Ly7. **J**, Genomic feature annotation of CREBBP-unique, KMT2D-unique, and CREBBP/KMT2D-co-bound ChIP-seq peaks. The definitions for different genomic features are depicted below the bar plot. **K**, Co-IP for assessing interaction between endogenous CREBBP and KMT2D in OCI-Ly7. CREBBP, KMT2D and IgG control were immunoprecipitated from OCI-Ly7 nuclear extracts. Western blots were performed using anti-CREBBP and KMT2D antibodies. **L-M**, ChIP-qPCR quantifying binding of endogenous (**L**) KMT2D and (**M**) CREBBP at the indicated gene loci in the indicated isogenic OCI-Ly7 cell lines. ChIP signals from three independent experiments were normalized to input and then to WT and presented as mean ± SD. The p values were calculated by unpaired t test and BH adjusted for multiple comparisons and denoted as follows: ns p > 0.05; *p < 0.05; **p < 0.01; ***p < 0.001; ****p < 0.0001. **N**, Functional annotation of C1 genes (n=401) by Enrichr and Toppgene (45, 46).

To examine the impact of these mutant genotypes on chromatin, we first conducted western blots for H3K27ac and H3K4me1 in our isogenic cells and observed no significant difference among the various clones and genotypes (**Supplementary Fig. S5I-K**). This did not preclude perturbation of the distribution of these marks throughout the genome, and therefore we also performed CUT&RUN (Cleavage Under Targets & Release Using Nuclease) profiling of H3K27ac, H3K4me1 and H3K4me3. For the unbiased analysis of these data we performed t-SNE dimensionality reduction and K-means clustering of the union of H3K27ac, H3K4me1, and H3K4me3 peaks from each genotype, which defined eight clusters with distinct patterns (**Fig. 6C**). As expected, the distribution of H3K27ac and H3K4me1 was highly overlapping in WT cells, although there were many H3K4me1 marked sites that lacked H3K27ac (also expected, potentially representing poised enhancers, **Fig. 6D**). In contrast there was very little overlap with H3K4me3. Comparing these with the perturbation induced by each mutant genotype revealed overlapping and progressive reduction in H3K27ac and H3K4me1 in C, K and CK isogenic cells that appeared strongest in the cluster 1 region (**Fig. 6C, 6E**). Shifts in H3K4me3 were milder and only partially overlapped with the other marks, although they were less affected in CK vs the other two mutant genotypes. Plotting mRNA abundance of genes corresponding to these loci (using GREAT) revealed progressive reduction of expression in genes mostly corresponding to the cluster 1 region (**Fig. 6E**). The unique properties of cluster 1 were further illustrated in a comparative cluster representation, showing that these loci are enriched for H3K27ac and H3K4me1 but not H3K4me3 in WT cells (**Fig. 6F**). Cluster 1 loci manifested progressive reduction of H3K27ac and H3K4me1 but not H3K4me3 in C, K and CK, the corresponding genes were progressively repressed, and cluster 1 loci were also distal to TSS, collectively suggesting that these regions corresponded mostly to gene enhancers (**Fig. 6F**). Adjacent clusters 2 and 3 appear as less affected versions of cluster 1, whereas clusters 4-6 represent gene promoters. Clusters 7-8 corresponded to genes strongly upregulated in all mutant genotypes, of unclear significance from the chromatin perspective. Cluster 1 loci/genes thus appear to be most directly dependent both on CREBBP and KMT2D.

Unexpectedly, analysis of cluster 1 loci showed that in addition to reduction of H3K27ac, loss of CREBBP resulted in reduction of H3K4me1 (**Fig. 6G-H, Supplementary Table 5**). Reciprocally, loss of KMT2D resulted in reduction of H3K27ac in addition to its effect on H3K4me1. This was more directly represented by plotting read density of cluster 1 peaks for each genotype, which further suggested that the effect of KMT2D loss was more severe on H3K27ac than the loss of CREBBP on H3K4me1(**Fig. 6G-H, Supplementary Table 5**). To better understand this phenomenon, we analyzed ChIP-seq for CREBBP and KMT2D in OCI-Ly7 cells (19, 20), which showed that 55% of KMT2D peaks overlap with CREBBP, whereas 38% of CREBBP peaks overlapped with KMT2D (**Fig. 6I**). Regardless of overlap, KMT2D and CREBBP peaks mapped mainly to intergenic, distal and intronic elements, consistent with their predominantly enhancer-related function (**Fig. 6J**). This interdependency led us to explore whether these two proteins could form a complex. We therefore performed endogenous protein co-immunoprecipitations in two independent GCB-DLBCL cell lines (OCI-Ly7 and SUDHL4), and observed reciprocal enrichment for each protein with their respective antibodies but not with control antibody (**Fig. 6K, Supplementary Fig. S5L**), with a higher fraction of KMT2D associated with CREBBP than vice versa, consistent with ChIP-seq analysis (**Fig. 6I**). We reasoned that such interaction might contribute to their stable chromatin association. To test this notion we performed ChIP quantitative PCR (ChIP-qPCR) assays for KMT2D in CREBBP knockout cells, and for CREBBP in KMT2D knockout cells, using primers corresponding to enhancers for genes that are particularly repressed in the CK setting, as well as negative control primers (**Fig. 6L-M**). We found that loss of CREBBP resulted in reduced enrichment for KMT2D (**Fig. 6L**), and vice versa loss of KMT2D resulted in reduced recruitment of CREBBP (**Fig. 6M**), arguing that formation of stable chromatin association is at least partially interdependent and perhaps linked to their propensity to form a complex together.

Finally, to better understand the transcriptional programs and biological functions linked to cluster 1 (i.e. the most CK dependent genes) we performed Enrichr and ToppGene analyses (45, 46). Taking together published ChIP-seq studies, motif analysis, and known TF networks, we observed significant enrichment for many of the CC immune synapse responsive TFs that drive GC exit cell fate decisions including SPI1, RELA, RELB, AP1, STAT3, IKZF1, BATF and others (padj<0.05, **Fig. 6N, Supplementary Table 2**). Accordingly, pathways and protein interaction networks were mostly immune synapse related including AP1, BCR, TLR, CTLA4 blockade, IFNg, CREBBP, SYK, BTK, DNA damage, etc. These were further associated with immune reaction phenotypes such as abnormal humoral immunity and B cell physiology, abnormal CD8 and CD4 responses, and others. We further validated that CK loss induced more severe repression than either loss alone of key immune synapse target genes in independent experiments at both the mRNA (RT-qPCR) and protein (FACS) levels (**Supplementary Fig. S5M-O**). In summary, these results indicate that CREBBP and KMT2D co-occupy inducible enhancers of immune synapse/GC exit cell fate determination pathways, where they are required for each other’s stable chromatin loading and hence histone modification activity, with their simultaneous deficiency eliciting a more profound perturbation of chromatin state of these enhancers than either single deficiency.

### Super-enhancers are particularly susceptible to CREBBP and KMT2D deficiency-induced perturbation

Cell fate determination is proposed to depend on super-enhancers to induce expression of phenotype-driving genes (47). Given that CK dependent enhancers were enriched for such genes and pathways, and were located quite distally to TSSs, we wondered whether the effect of their co-depletion might disproportionately affect super-enhancers. Using our murine GC B cell ATAC-seq data we compared and contrasted differential accessibility at enhancers and super-enhancers. Whereas effects on standard enhancers were fairly small and similar among mutant genotypes, super-enhancers were clearly more severely affected, especially in the K and CK settings, with CK being even further significantly impaired than K (see 95% confidence intervals, **Fig. 7A**). We then tabulated differential accessibility across constituent peaks for each super-enhancer and ranked them based on their log2 fold change in CK vs WT. We linked super-enhancers to genes within the same mouse GC topologically associated domains (TADs) using GREAT (48). Importantly, these included many immune synapse response and GC exit genes, as exemplified by Cd86, Il9r and Zbtb20 (**Fig. 7B**), which are most strongly downregulated in CK-deficient CB and CC (**Fig. 3**).

**Figure 7.**
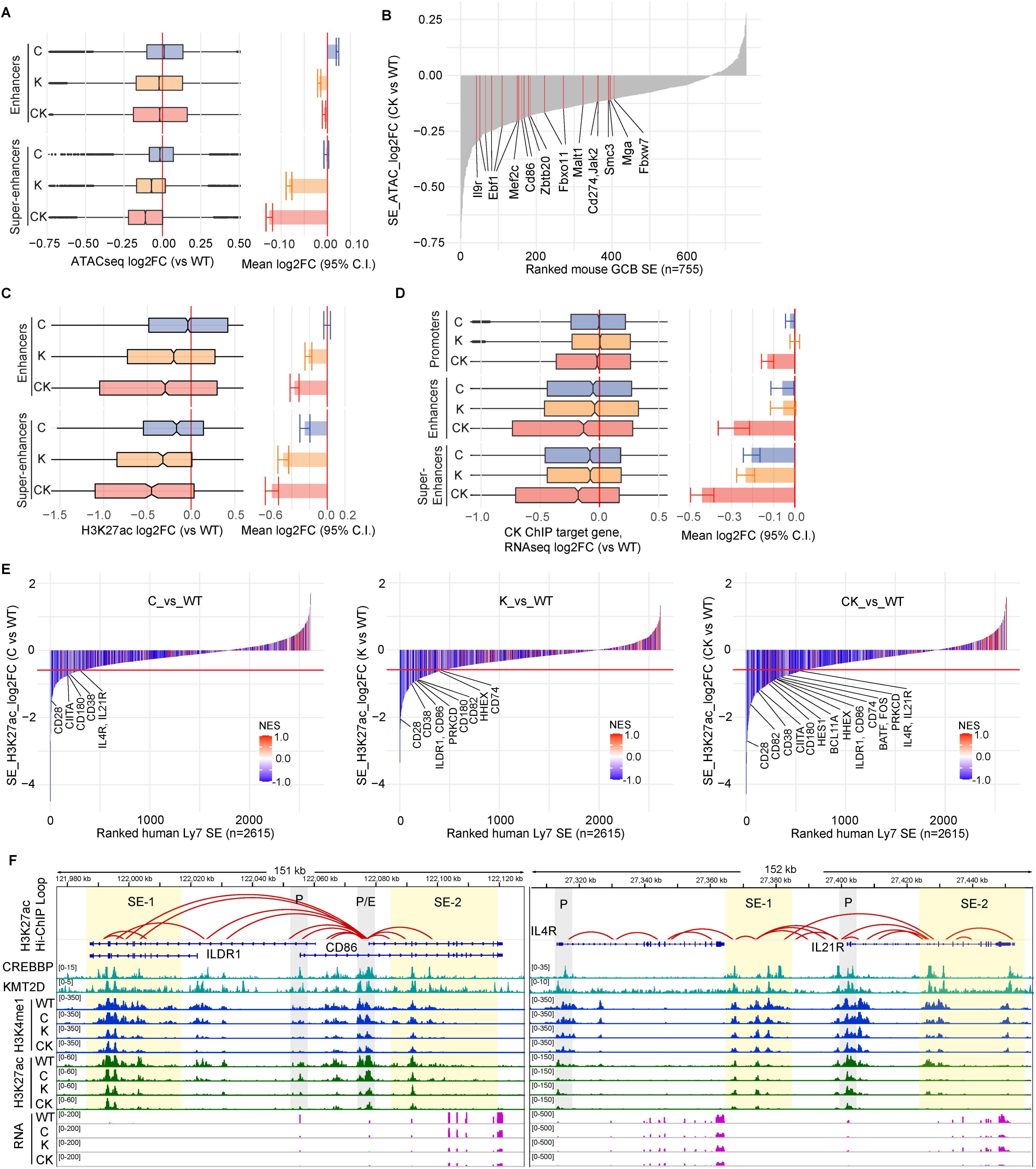
Crebbp and Kmt2d haploinsufficiency cooperatively and preferentially perturbs activation of super-enhancers critical for GC cell fate specification. **A**, Left: box plots showing chromatin accessibility change at enhancer and constituent super-enhancer peaks in C, K and CK vs WT mouse GC B cells. The line in the middle of the box marks the median. The vertical size of the box denotes the interquartile range (IQR). The upper and lower hinges correspond to the 25th and 75th percentiles. Right: bar plots showing mean of the corresponding chromatin accessibility log2FC (C/K/CK vs WT), with error bars indicating 95% confidence intervals (C.I.) of the mean. **B**, Waterfall plot ranking super-enhancers (SE) in mouse GC B cells based on their accessibility change in CK vs WT. Constituent ATAC-seq peaks in each SE were summed before calculating the fold change. Red-highlighted closing SEs are linked by GREAT to the labeled genes downregulated in CK_CC vs WT_CC, and located within the same mouse GC defined TADs. **C**, Left: box plots showing H3K27ac signal change at enhancer and constituent super-enhancer peaks in C, K and CK vs WT OCI-Ly7 cells. The line in the middle of the box marks the median. The vertical size of the box denotes the interquartile range (IQR). The upper and lower hinges correspond to the 25th and 75th percentiles. Right: bar plots showing mean of the corresponding H3K27ac log2FC (C/K/CK vs WT), with error bars indicating 95% confidence intervals of the mean. **D**, Left: box plots showing the C/K/CK deficiency-induced expression changes in the target genes of CK-co-bound promoters, enhancers, and super-enhancers in OCI-Ly7 cells. Right: bar plots showing mean of the corresponding RNA log2FC (C/K/CK vs WT), with error bars indicating 95% confidence intervals of the mean. **E**, Waterfall plots ranking super-enhancers (SE) in OCI-Ly7 cells based on their H3K27ac level change in C (left), K (middle) or CK (right) vs WT. Constituent H3K27ac peaks in each SE were summed before calculating the fold change. Bars are colored by normalized enrichment scores (NES) of RNA-seq GSEA, which used all genes in each SE-residing TADs as gene signature against ranked gene list based on their expression change in C/K/CK vs WT. Highlighted genes are downregulated in C/K/CK vs WT with log2FC < −0.58 (red line) and linked by GREAT to the closing SEs located within the same human GC defined TADs. **F**, IGV views of normalized histone marks CUT&RUN and RNA-seq signals at the indicated loci in isogenic OCI-Ly7 cells. H3K27ac Hi-ChIP loop calls are depicted as arcs connecting the two interacting loci. Promoters and super-enhancers are shaded in gray and yellow, respectively.

Examining the impact of these three mutant genotypes on H3K27ac distribution in isogenic human cells again showed that reduction of this histone mark is more severe at the constituent peaks of super-enhancers than regular enhancers, with the fold change loss in CK super-enhancers being most dramatic (**Fig. 7C**). To more specifically compare and contrast the impact of CK deficiency against promoters, enhancers or super-enhancers we focused on such regulatory elements with ChIP-seq confirmed binding of both CREBBP and KMT2D, and examined their impact on gene expression. Most notably, target genes of CK-bound super-enhancers were more repressed than those of CK-bound regular enhancers upon C, K or CK loss, with the latter reducing the target gene expression to a substantially greater extent than either single loss alone (**Fig. 7D**). None of the mutant genotypes had much impact on gene expression based on promoter CK binding. We next summed the H3K27ac signal of constituent peaks for each super-enhancer, and ranked them based on log2 fold change between C/K/CK and WT, and linked them to their corresponding genes within the same human GC TADs (49) (**Fig. 7E**). To further confirm that super-enhancers of interest interact with their respective genes, we examined their connectivity based on available H3K27ac HiChIP performed in OCI-Ly7 cells (50). Plotting the impact of C, K or CK deficiency on super-enhancers that connect with important immune synapse responsive genes such as *CD86* and *IL21R* further allowed direct visualization of the progressive loss of H3K27ac, H3K4me1 on relevant super-enhancers and the corresponding suppression of gene expression (**Fig. 7F**). These data suggest that the cooperative effect of CREBBP and KMT2D loss of function on B cell phenotype and lymphomagenesis may be due to a more critical biochemical need for CK complex formation at distal immune-synapse-responsive super-enhancers linked to GC exit and cell fate decisions in the GC reaction, vs regular enhancers that are normally located more proximal to TSSs.

### CREBBP and KMT2D deficiency suppresses a core immune signature in GC B cells that is retained and shared between murine and human lymphomas

Results described above point to a cooperative disruption of immune-synapse responsive genes upon CK loss observed both in GC B cells and established human DLBCL cell lines. We reasoned that CK gene expression signatures induced by CK deficiency in GC and persisting upon malignant transformation would indicate these effects most likely provide a selective advantage to lymphoma cells. For this, we first performed RNA-seq profiling of sorted B220^+^ lymphoma cells from each genotype at day 235 of experiment shown in **Fig. 1**. We then identified the set of genes repressed in BCL2+CK relative to BCL2 mouse tumors. These genes were applied to a GSVA-based estimation of their relative expression across an RNA-seq dataset derived from human FL patients (n=16,10,8,15 for epigenetic WT, C, K, and CK, respectively) (19). This analysis indicated that the CK repressed gene signature was progressively decreased from C, K, to CK compared to patients without C, K or *EZH2* mutations (defined as epigenetic WT), with the decrease being significant only for the CK cases (**Fig. 8A**). We then performed a comparative cross-species analysis of CK repressed signatures in murine CBs, CCs and lymphoma cells, as well as human DLBCL cell lines and human FL patients, which identified a list of CK target genes that exhibit consistent downregulation compared to WT in at least two different datasets (**Fig. 8B, Supplementary Table 2**). Once again this list contained numerous critical immune synapse-response genes such as *CD86*, *BTLA*, *TLR4*, etc. Functional annotation of this CK_consistent_down gene set using Enrichr recapitulated our findings in human DLBCL cell lines, with disruption of GC exit TFs based on ChIP-seq, motif or network analysis including BATF, IKZF1, AP1, SPI1, RELB, SPIB, HHEX, etc (**Fig. 8C**). Functional annotation of this gene set enriched for BCR, NFkB, IFNg, chemokine, CD8 and other CB related genes such as ATR and G2/M checkpoint pathways; and phenotypes such as abnormal response to antigen and abnormal B cell activation. These data point to sustained suppression of immune synapse genes as contributing to transformation and maintenance of CK mutant lymphomas.

**Figure 8.**
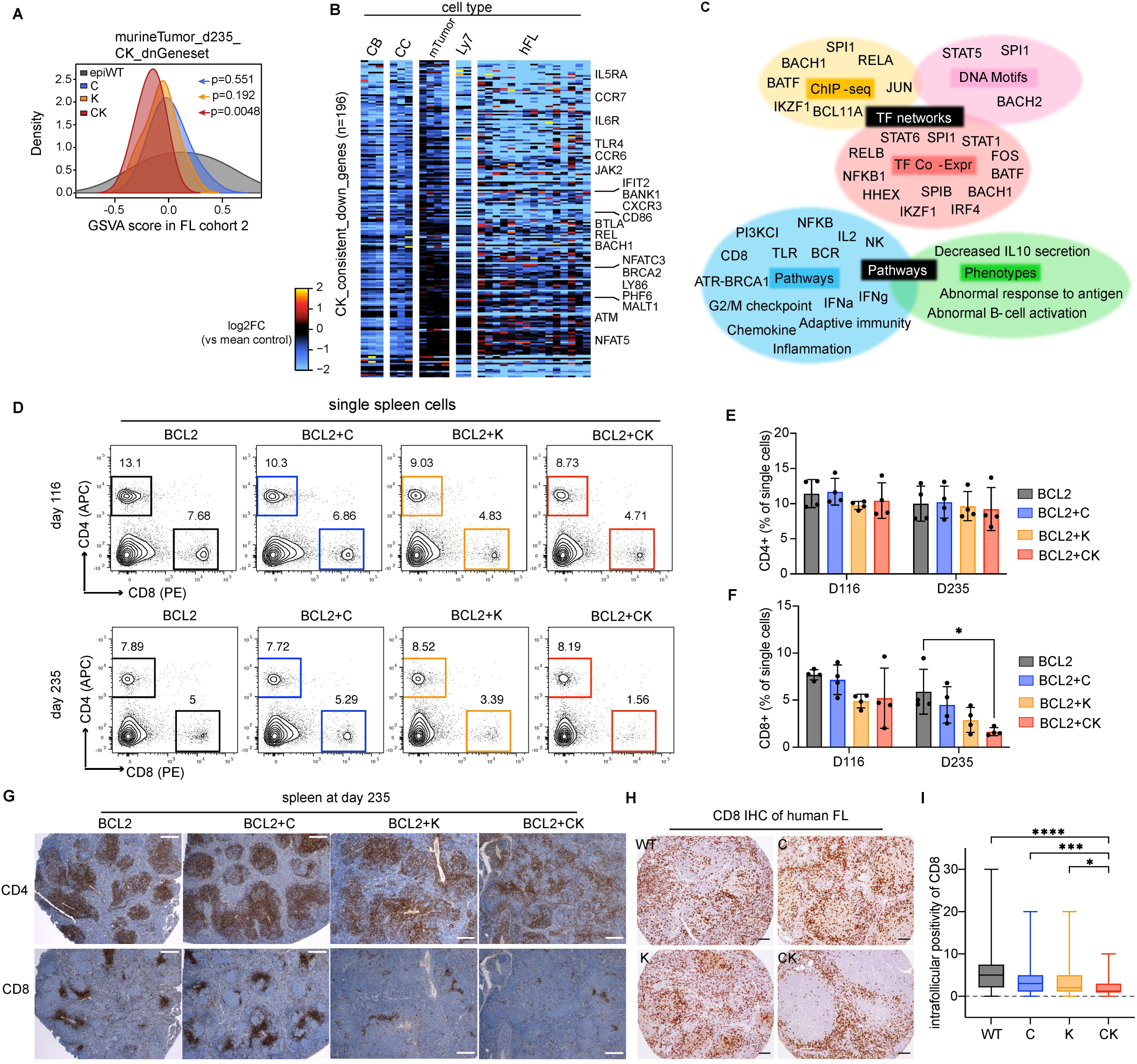
CK deficiency-induced murine lymphomas recapitulate the transcriptional perturbation and low CD8+ T cell infiltration in CK mutant human lymphomas. **A**, GSVA analysis using genes downregulated in BCL2+CK vs BCL2 murine lymphoma cells at day 235 as gene set against RNA-seq datasets of human FL_cohort2 patients stratified based on C and K mutation status (n=16,10,8,15 for epiWT, C, K, and CK respectively). The p values were calculated using two-sided Wilcoxon test. **B**, Heatmap showing the relative expression levels of CK_consistent_down_genes (n=196), which include genes exhibiting significant downregulation in at least two of the indicated RNA-seq datasets and downregulated but not significantly in the remaining datasets. **C**, Functional annotation of CK_consistent_down_genes by Enrichr and Toppgene. **D**, Representative FACS plots show the gating strategy and relative frequency of splenic CD4^+^ and CD8^+^ T cells in mice from experiment shown in Figure 1 at days 116 and 235 post-BMT. **E-F**, FACS analysis showing the relative abundance of splenic (**E**) CD4^+^ T cells and (**F**) CD8^+^ T cells normalized to total single cells at days 116 and 235 post-BMT (mean ± SD). Each dot represents a mouse (n=4 mice per genotype). Statistical significance was determined using ordinary one-way ANOVA followed by Tukey-Kramer’s multiple comparisons test (*p < 0.05). **G**, Representative CD4 and CD8 IHC staining images of spleen sections from mice euthanized at day 235 post-BMT. The scale bar represents 200 pixels. **H**, Representative CD8 IHC staining images of human FL tissue microarrays (TMAs) (51). The scale bar represents 100 pixels. **I**, Box plot showing the relative percentage of CD8^+^ T cells among overall cellularity in the human FL TMAs as shown in panel **H** (n=33, 43, 47, and 88 for WT, C, K, and CK human FLs respectively). The p values were calculated by Kruskal-Wallis test followed by Dunn’s multiple comparisons test (*p < 0.05, ***p < 0.001, ****p < 0.0001).

### CREBBP+KMT2D mutant lymphomas manifest greater depletion of CD8^+^ T cells

This consistent repression of B cell to T cell immune synapse genes led us to consider whether there was any potential consequence of such findings for the immune microenvironment of CK lymphomas. We therefore assessed the abundance of CD4^+^ and CD8^+^ T cells by flow cytometry from early and late stage BCL2, BCL2+C, BCL2+K and BCL2+CK lymphomas from the murine model shown in **Fig. 1**. These studies showed similar abundance of CD4^+^ cells across all genotypes at both time points (**Fig. 8D-E**). In contrast, there were progressively lower numbers of CD8^+^ cells that were only statistically significant in the BCL2+CK setting (**Fig. 8D, 8F**). This was only obvious at the later time point, although a mild trend was apparent in the earlier lymphomas. We further explored this finding by performing CD4 and CD8 immunohistochemistry staining on late-stage lymphomas from all four genotypes. We found that CD4^+^ cells were approximately similar in abundance across all cases (**Fig. 8G**). However, whereas CD4^+^ cells were mostly confined to malignant follicles in BCL2 mice, their distribution became progressively more disseminated across the BCL2+C, BCL2+K and BCL2+CK lymphomas. CD4^+^ follicular clustering was severely disrupted in BCL2+K and completely lost in BCL2+CK mice. Strikingly, there was a clear reduction in CD8^+^ cell abundance that was most profound in BCL2+CK mice (**Fig. 8G**). In contrast to CD4^+^ cells, CD8^+^ cells were mostly localized to perivascular spaces, as expected. Although blood vessels were clearly present in BCL2+CK mice, very few CD8^+^ T cells were able to accumulate at these sites (**Fig. 8G**). To determine whether a similar phenomenon might occur in humans we examined CD8 staining patterns using a tissue microarray representing 211 FL patient specimens, that were also characterized by mutation profiling (51). These studies showed significant exclusion of CD8^+^ cells from malignant follicles in CK mutant human FLs as compared to C, K or epigenetic WT patients (**Fig. 8H-I**). Together, these data suggest that more potent, CK-induced loss of signaling receptors that mediate bi-directional crosstalk between B and T cells, such as the CD28 costimulatory ligand CD86 protein, may lead to more severe disruption of immune-surveillance and hence a stronger selective advantage for cells with these mutations.

## DISCUSSION

Herein, we explored the puzzling co-occurrence of somatic mutations in the two enhancer activating chromatin modifier proteins CREBBP and KMT2D in B-cell lymphomas. We confirmed that these mutations are genetically highly co-occurrent in additional FL and GCB-DLBCL cohorts. We found that conditional heterozygous knockout of these chromatin modifiers does indeed cooperate to yield a more aggressive lymphoma phenotype, suggesting that this combination is especially beneficial in providing a selective advantage to pre-malignant B cells. We did not model double homozygous loss of function, since this scenario rarely if ever occurs in humans, and we speculate they may confer an unfavorable functional state to GC B cells. It remains unknown if there is a specific order that these mutations need to occur in the pre-lymphoma state, since presumed clonal precursor B cells have been identified with both of these genetic lesions. *CREBBP* HAT mutations were shown to have similar, but more potent loss of function effects than heterozygous deletion (52). The fact that we observed similar cooperative epigenetic and transcriptional perturbations in our heterozygous mouse model as in *CREBBP*^R1446C^ DLBCL cells further supports this notion.

It is well-established that H3K4me1 and H3K27ac prime and activate enhancers, respectively. In contrast to this sequential action model, our data support a reciprocal cooperation model. Specifically, we show that CREBBP interacts with KMT2D, they are reciprocally required for each other’s stable chromatin association, and that losing either enzyme reduces the levels of both H3K4me1 and H3K27ac. This reciprocal cooperation model is plausible given how these histone marks are normally regulated during the GC reaction. Specifically, previous reports showed that enhancers activated by CREBBP and KMT2D become transiently repressed in GC centroblasts. This was due to actions of BCL6, a GC master regulatory transcriptional repressor. BCL6 binds mainly to enhancers, where it mediates H3K27 deacetylation through recruitment of HDAC3, and H3K4me1 demethylation through recruitment of KDM1A (19,53,54). It is believed that BCL6 repressor complexes dissociate in the LZ, allowing CREBBP and KMT2D to rapidly and simultaneously restore enhancer activation marks (55, 56), activities of which are likely further enhanced through immune synapse signals received during T cell help.

Given these considerations, our data suggest that the non-catalytic scaffold function of CREBBP and KMT2D plays an important role in their intimate cooperation, reminiscent of a prior study reporting that UTX bridges the interaction between P300 and KMT2D (57). However, there are other potential ways that CREBBP and KMT2D could tune each other’s activity. For example, it is possible that CREBBP-catalyzed H3K27ac weakens histone tail-DNA interactions to open chromatin structure, which in turn makes the H3K4 site accessible to KMT2D (58). Conversely, KMT2D may promote CREBBP-catalyzed H3K27ac indirectly through downstream effects of H3K4me1, which was reported to recruit chromatin remodeler complex BAF, cohesin complex and antagonize DNA methylation (59–62). Furthermore, KMT2D was recently appreciated to contain intrinsically disordered regions (IDRs) that confer its ability to undergo liquid-liquid phase separation (63). Interaction of CREBBP with KMT2D could help to concentrate CREBBP into the phase separated transcriptional condensates, which enhance the enzymatic activity of CREBBP in depositing histone acetylation (64). We also cannot rule out that some portion of the observed effects could be linked to putative roles of these proteins on non-histone proteins. For example, it is possible that CREBBP and KMT2D could induce post-translational modifications in each other (65), which could modulate their activity. Finally, it is likely that the presence of the TFs that we identified as enriched at KMT2D and CREBBP target sites could provide further stability to the interaction of these proteins. Indeed a recent study showed that PRC1 complex required the presence of both PRC2 and the transcription factor BCL6 for stable association with chromatin in GC B cells (66). Teasing apart the relative contribution of these possible mechanisms will require an extensive biochemical effort and will provide additional important basic insight into the effects we observed herein.

It is notable that super-enhancers were especially vulnerable to depletion of both CREBBP and KMT2D, which may offer a key explanation for why their mutations are co-occurrent. Super-enhancers are complex regulatory elements composed of multiple clustered enhancers, often quite distal from the genes they regulate, and have been described as regulating genes required for cell fate decisions (47). Typically, super-enhancers contain binding sites for many TFs, which recruit many different chromatin modifying proteins (64). Although some reports claim that clustered enhancers can act in a synergistic manner, it is clear that many super-enhancers are more additive in nature. This is consistent with our finding of additive loss of super-enhancer function induced by loss of CREBBP and KMT2D alleles. A super-enhancer effect may also be inferred by the additive cell fate decision defect observed, whereby CCs were progressively impaired in their ability to exit the GC reaction as plasma or memory cells from C to K to CK mice. The transcriptional profile observed suggests persistence of CB transcriptional features in CCs, which may favor their retention in the DZ rather than exit.

It is notable that many of these CK dependent super-enhancer regulated genes encode for immune synapse signaling proteins that mediate cross talk with T cells, and direct cell fate upon T cell help and GC exit. The hallmark of GCB-DLBCL and FL is persistence of the GC transcriptional program that includes transient BCL6-mediated downregulation of many immune receptors that become re-expressed in the LZ. Failure of these genes to re-activate is thus central to the phenotype of these tumors and may explain why *CREBBP* and *KMT2D* mutations are so highly prevalent, given that we show how strongly they impair these programs. Indeed C+K deficiency impaired gene activation by most of the canonical LZ TFs including SPI1, BATF, NFkB, STAT3, AP1, IKZF1 and others (41, 42). This manifested as impaired expression of key genes involved in GC B cell interaction with TFH cells and GC exit. Notably among these was robust repression of CD86, an important T cell costimulatory ligand that binds to CD28 (67), as well as other key co-stimulatory genes such as CD40 and SLAMF1 (68, 69), and GC regulating chemokine receptors such as CCR6 (70), the absence of which enhances GC formation and impairs affinity maturation. This was coupled with stronger silencing of MHC class II than what was observed for example with CREBBP haploinsufficiency alone. Collectively it might be anticipated that the sum of these perturbations would lead to reduction in immune tone and hence impaired activation and recruitment of T cells.

These considerations may play into the observed reduction in CD8^+^ T cells that was especially strong in CK murine and human lymphomas. The impaired ability of CD8^+^ cells to engage with CK deficient lymphoma may represent a form of aberrant super-enhancer mediated immune evasion, which may contribute to accelerated pathogenesis, and have implications for the relative efficacy of T cell directed immunotherapies. Along these lines it was shown that HDAC3 selective inhibitors given to counteract HDAC3 antagonism of CREBBP, recruited T cells into murine GCB-DLBCLs in vivo and enhanced the anti-tumor efficacy of checkpoint blockade (52). Moreover, CREBBP mutant lymphoma cells were also more sensitive to HDAC3i in a cell autonomous manner, which has led to the concept of using these drugs for precision therapy in FL and DLBCL. Conversely, it was recently shown that *KMT2D* mutation sensitizes lymphoma cells to KDM5 inhibitors (71). KDM5 histone demethylases remove H3K4me3 rather than H3K4me1, hence these drugs were thought to mediate their effects by impairing dynamic changes in H3K4 methylated states. We would predict that CK lymphomas would be highly sensitive to these agents given the cooperative actions of these two corepressors, and perhaps even more to their combination. Hence our findings provide a basis for development of epigenetic combinatorial therapy with these agents, to reverse the effects of these mutations and induce both cell autonomous and immune anti-tumor activity. As these drugs become available, we propose they be tested in this context and used as immune adjuvant therapy together with T cell enhancing immunotherapies.

---

**Supplementary Figure 1.**
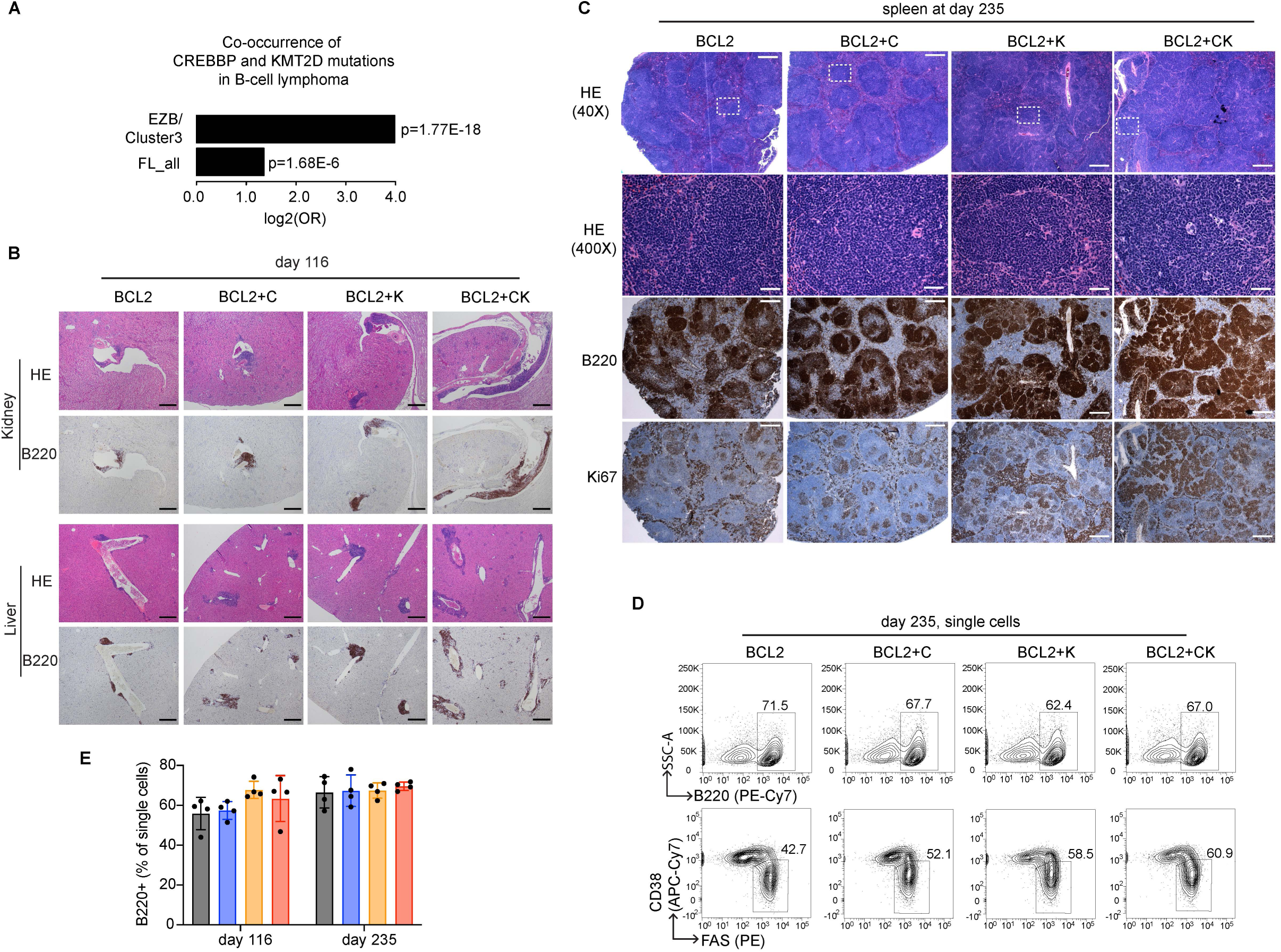
**A**, Odds ratio (OR) analysis of *CREBBP* and *KMT2D* mutations among EZB/Cluster3 DLBCL (n=319) and FL_all (merged FL datasets, n=478) cohorts of patients. The p values were calculated by Fisher’s exact test. **B**, Representative H&E and B220 IHC images of formalin-fixed paraffin-embedded kidney and liver sections prepared from mice euthanized at day 116 post-BMT. The scale bars represent 380 pixels. **C**, Representative H&E, B220 and Ki67 IHC images of formalin-fixed paraffin-embedded spleen sections from mice euthanized at day 235 post-BMT. The scale bars represent 200 pixels. **D**, Representative FACS plots show the gating strategy and frequency of B220^+^CD38^-^ FAS^+^ splenic GC B cells in mice at day 235 post-BMT. **E**, FACS analysis showing the relative abundance of splenic total B cells (B220^+^) normalized to total single cells at day 116 and 235 post-BMT (mean ± SD). Each dot represents a mouse (n=4 mice per genotype).

**Supplementary Figure 2.**
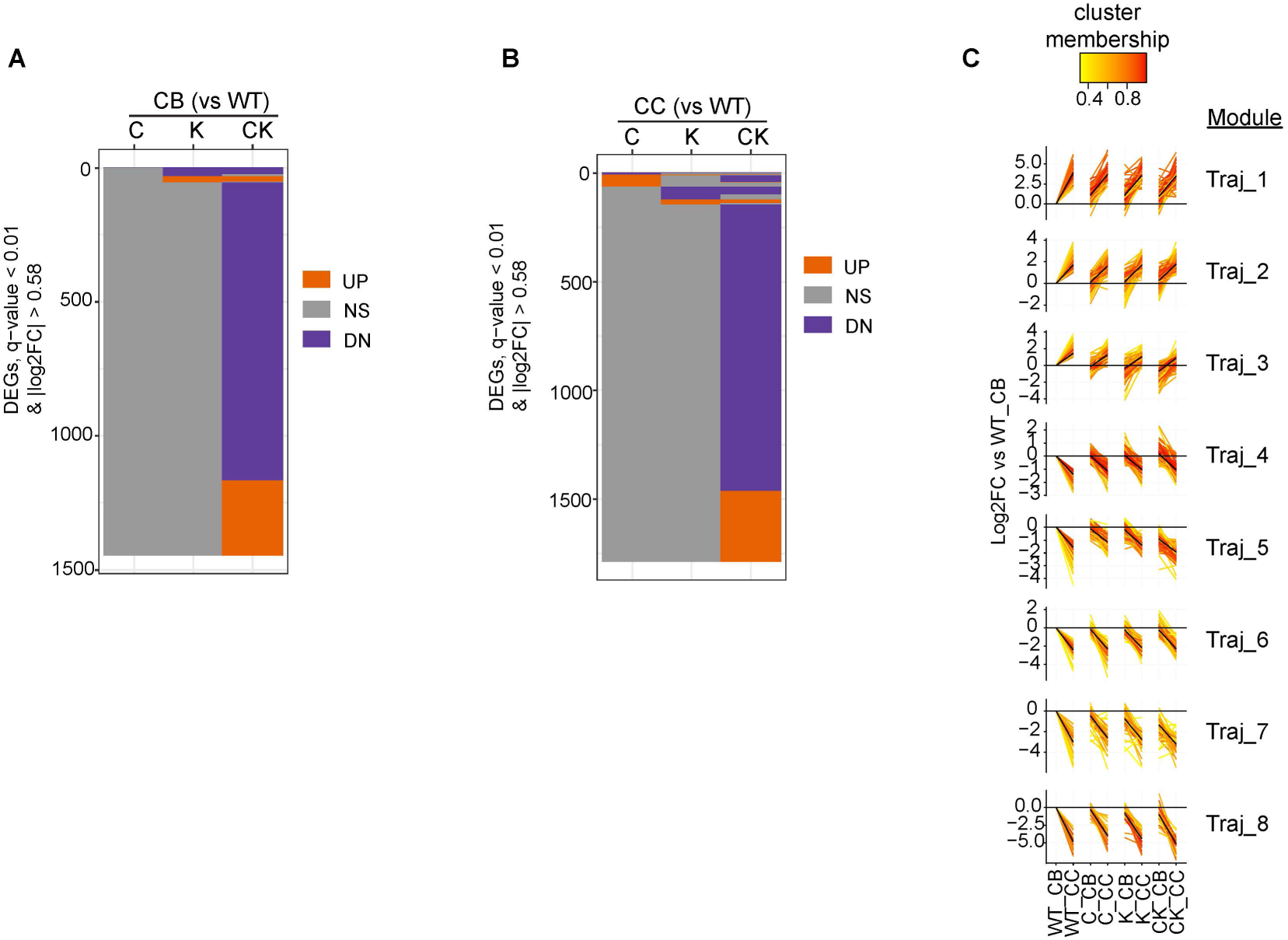
**A-B**, Bar plot comparing the gene expression status and showing DEG overlap in (**A**) CB and (**B**) CC among C, K, and CK relative to WT. Purple represents downregulated, orange depicts upregulated and gray represents no differentially expressed genes as compared to WT (|log2FC| > 0.58, q < 0.01). **C**, Fuzzy c-means clustering of RNA-seq datasets identified 8 clusters (named as Traj_1 to Traj_8) with distinct trajectory pattern: line plot of standardized log2 fold-change relative to mean of WT CB. Black lines in the line plot are cluster centroid; each gene is colored by the degree of cluster membership.

**Supplementary Figure 3.**
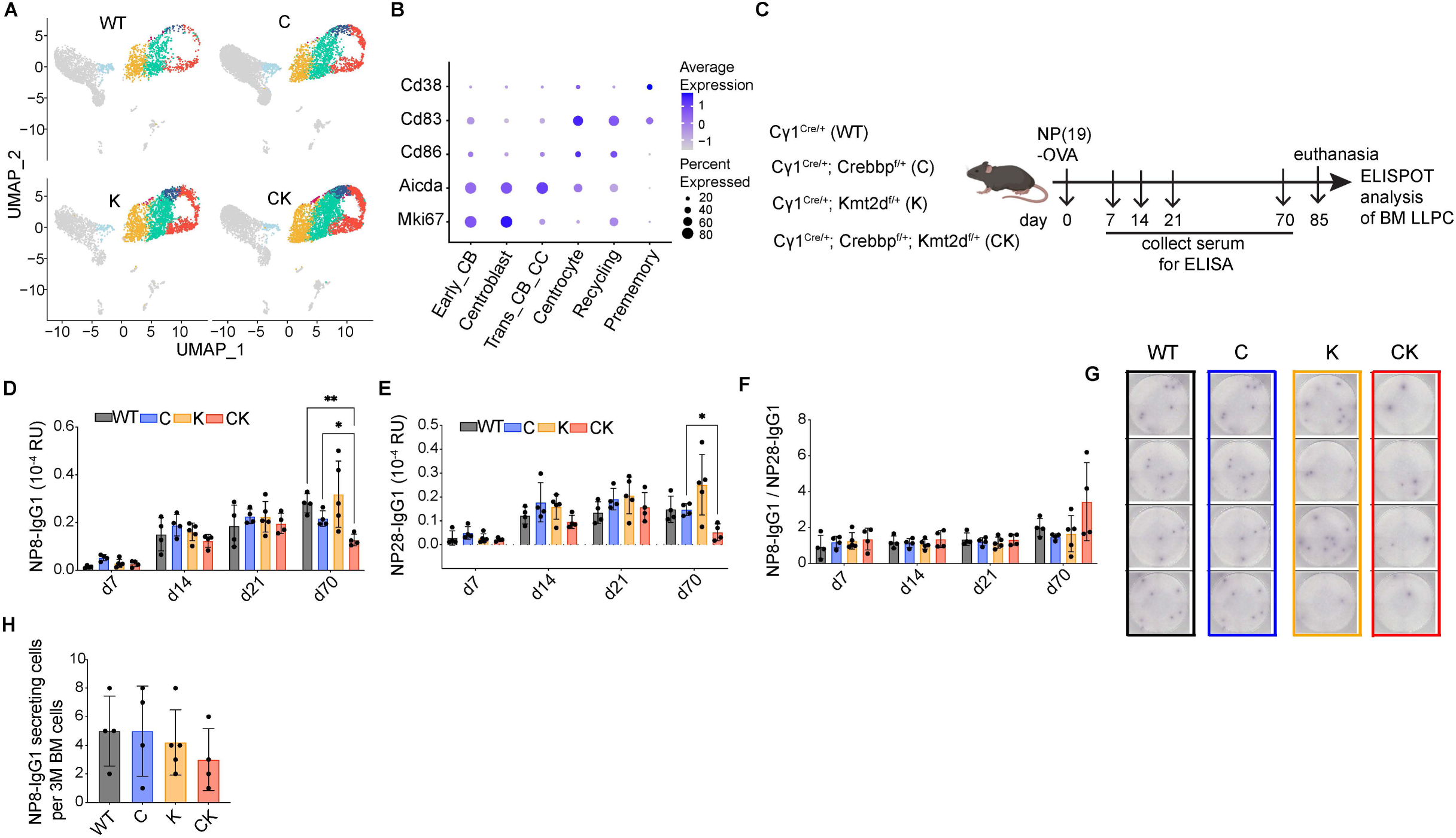
**A**, UMAP representation of single-cell RNA-seq datasets for each genotype. **B**, Dot plot showing the average expression levels of the indicated genes and percent of cells expressing the genes in each GC B cell subpopulation. **C**, Experimental design for (**D-H**). **D-F**, ELISA analysis showing (**D-E**) the titers and (**F**) ratios of NP-specific high- (**D**, NP8-reacting) or low-affinity (**E**, NP28-reacting) IgG1 in serum collected at the indicated time points post NP-OVA immunization (mean ± SD). Each dot represents a mouse. Statistical significance was determined using two-tailed unpaired t test (*p < 0.05; **p < 0.01). **G**, Representative ELISPOT images for each genotype. Each spot represents a long-lived, plasma cell (LLPC) in the bone marrow capable of secreting NP-specific high-affinity IgG1 antibody (NP8+ IgG1). **H**, ELISPOT assay showing the number of NP8-IgG1-secreting LLPCs per 3 million total BM cells for each genotype (mean ± SD). Each dot represents a mouse.

**Supplementary Figure 4.**
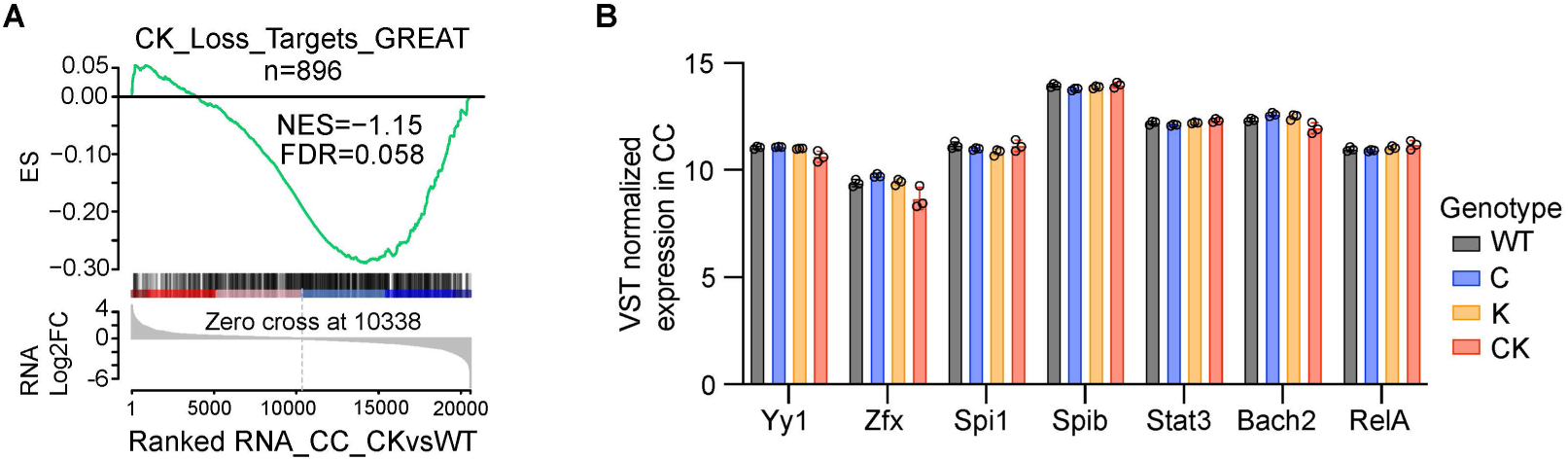
**A**, GSEA analysis using CK_Loss ATAC-seq peaks-targeting genes (annotated by GREAT) as the gene set against a ranked gene list based on CK_CC vs WT_CC RNA-seq. NES, normalized enrichment score. The p value was calculated by an empirical phenotype-based permutation test. The FDR is adjusted for gene set size and multiple hypotheses testing. **B**, Bar plot showing VST normalized expression values of the indicated TFs in mouse CC for each genotype as mean ± SD.

**Supplementary Figure 5.**
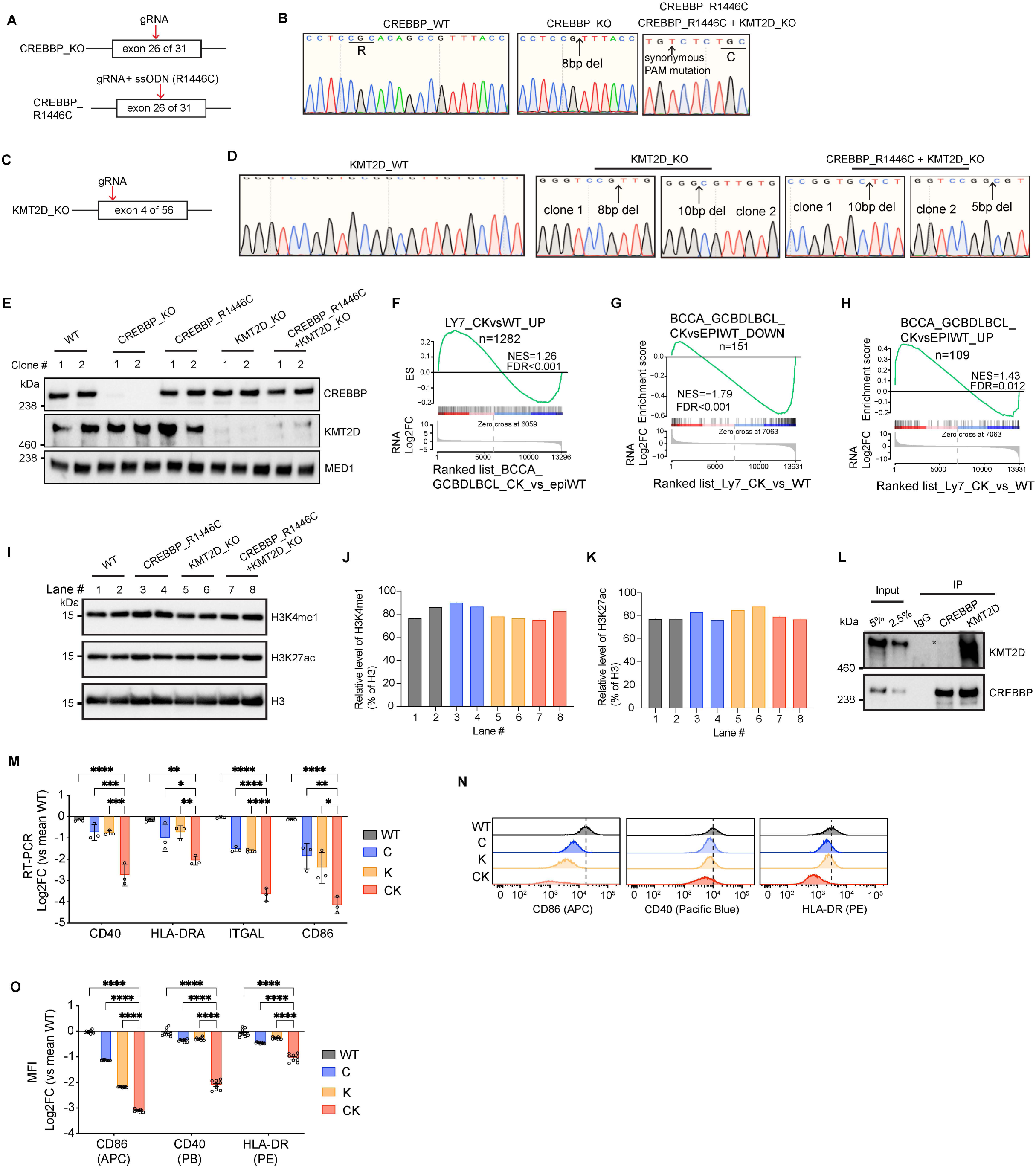
**A**, Experimental design for generating *CREBBP*-KO and *CREBBP*-R1446C, an enzymatically dead mutant, OCI-Ly7 cell lines. ssODN: single-stranded oligodeoxynucleotide. **B**, Sanger sequencing chromatogram confirming the homozygous knock out or R1446C point mutation in *CREBBP*. **C**, Experimental design for generating *KMT2D*-KO OCI-Ly7 cell line. **D**, Sanger sequencing chromatogram confirming the homozygous knock out of *KMT2D*. **E**, Immunoblotting for endogenous CREBBP and KMT2D in the indicated isogenic OCI-Ly7 cell lines using MED1 as internal loading control. **F**, GSEA plot using CK vs WT upregulated genes in OCI-Ly7 as the gene set against a ranked gene list based on CK vs epigenetic WT RNA-seq datasets of human BCCA cohort GCB-DLBCL patients. NES, normalized enrichment score. The p value was calculated by an empirical phenotype-based permutation test. The FDR is adjusted for gene set size and multiple hypotheses testing. **G-H**, GSEA plots using CK vs epigenetic WT (**G**) downregulated or (**H**) upregulated genes in human BCCA cohort GCB-DLBCL patients as the gene set against a ranked gene list based on CK vs WT OCI-Ly7 RNA-seq datasets. NES, normalized enrichment score. The p value was calculated by an empirical phenotype-based permutation test. The FDR is adjusted for gene set size and multiple hypotheses testing. **I**, Immunoblot for H3K4me1, H3K27ac and H3 in isogenic OCI-Ly7 cells. **J-K**, Relative densitometry of (**J**) H3K4me1 and (**K**) H3K27ac for panel **I**. **L**, Co-IP for assessing interaction between endogenous CREBBP and KMT2D in human SUDHL4 GCB-DLBCL cell line. **M**, RT-qPCR of indicated genes in different isogenic OCI-Ly7 cells. qPCR signal for each gene was normalized to those of HPRT and then mean WT and presented as log2 fold-change ± SEM. Statistical significance was determined using ordinary one-way ANOVA followed by Tukey-Kramer’s multiple comparisons test (**p < 0.01, ***p < 0.001, ****p < 0.0001). **N**, Stacked flow cytometry histograms showing the progressive signal decrease for the indicated surface markers in C, K, and CK-deficient OCI-Ly7 cells compared to WT. **O**, FACS measuring cell surface levels of the indicated markers in different isogenic OCI-Ly7 cells. Mean fluorescence intensity (MFI) of each surface marker was normalized to mean WT and presented as log2 fold-change ± SEM. Each dot represents a biological replicate. Statistical significance was determined using ordinary one-way ANOVA followed by Tukey-Kramer’s multiple comparisons test (****p < 0.0001).

## Methods

### Mouse Models

Animal care was in strict compliance with institutional guidelines established by the Weill Cornell Medical College, the Guide for the Care and Use of Laboratory Animals (National Academy of Sciences 1996), and the Association for Assessment and Accreditation of Laboratory Animal Care International.

The following strains were obtained from The Jackson Laboratory (Bar Harbor, ME, USA): C57BL/6J (CD45.2/2, stock 000664, RRID:IMSR_JAX:000664), B6.SJL-Ptprca Pepcb/BoyJ (CD45.1/1, stock 002014, RRID:IMSR_JAX:002014), Cg1-Cre (stock 010611, RRID:IMSR_JAX:010611), Crebbp^flox^ (stock 025178, RRID:IMSR_JAX:025178). Kmt2d^flox^ (RRID:IMSR_JAX:032152) mice were obtained from Kai Ge at NIDDK (33). The VavP-BCL2 (RRID:MGI:3842939) (35) model was developed by J.M. Adams (Walter and Eliza Hall Institute of Medical Research, Australia).

### Cell Lines

The human GCB-DLBCL cell line OCI-Ly7 (RRID:CVCL_1881) was obtained from Ontario Cancer Institute and cultured in IMDM (ThermoFisher; 12440061) supplemented with 15% FBS, 1% L-Glutamine and 1% penicillin-streptomycin. The human GCB-DLBCL cell line SU-DHL4 (RRID:CVCL_0539) was obtained from DSMZ and grown in RPMI 1640 (ThermoFisher; 11875093) with 10% FBS, 1% L-Glutamine and 1% penicillin-streptomycin. Cell lines were authenticated by Biosynthesis using their STR Profiling and Comparison Analysis Service, and are routinely tested for mycoplasma contamination in the laboratory.

### Germinal Center (GC) Assessment in Mice

To induce GC formation, age- and sex-matched mice were immunized intraperitoneally at 8 to 12 weeks of age with either 0.5 ml of 1:10 diluted sheep red blood cell (SRBC) suspension in PBS (Cocalico Biologicals) or 100 ug of the highly substituted hapten NP (NP_16_ to NP_32_) conjugated to the carrier protein ovalbumin (OVA) (Biosearch Technologies; N-5051) absorbed to Imject Alum Adjuvant (ThermoFisher; 77161) at a 1:1 volume ratio.

S-phase percentage of GC B cells was determined by EdU incorporation assay. Specifically, each mouse received 1mg EdU dissolved in PBS through retro-orbital injection 1 hour before euthanasia. After surface marker staining, splenocytes were subjected to fixation, permeabilization, and click reaction to fluorescently label incorporated EdU with AF488 following the instruction from the Click-iT Plus EdU Flow Cytometry Assay Kit (ThermoFisher; C10632).

### Bone Marrow Transplantation and Lymphomagenesis

Bone marrow (BM) cells were harvested from the tibia and femur of 8-12 weeks old donor mice. After red blood cell lysis using the ACK lysing buffer (Lonza; BP10-548E), 1 x 10^6^ cells were injected into the retro-orbital sinus of lethally irradiated C57BL/6J recipient mice (2 doses of 450 rad, 12 hours apart, on a Rad Source Technologies RS 2000 X-ray Irradiator). 8 weeks after transplantation to ensure full engraftment, mice were immunized with SRBC periodically (once every 3 weeks) to induce GC formation and the cells of origin for lymphomagenesis. Except for those euthanized at two specific time points (day 116 and day 235), all remaining mice were monitored until any one of several euthanizing criteria were met, including severe lethargy, more than 10% body weight loss, and palpable splenomegaly that extended across the midline, in accordance with Weill Cornell Medicine Institutional Animal Care and Use Committee–approved animal protocol (protocol #2011-0031).

To generate mixed BM chimeric mice, B6.SJL mice were lethally irradiated with two doses of 450 rad X-ray 12 hours apart and then transplanted with 1 x 10^6^ mixed BM cells containing equal ratio of WT (CD45.1/2) and CK (CD45.2/2) through retro-orbital injection.

### Flow Cytometry Analysis and Cell Sorting

Single-cell suspension of mononuclear mouse splenocytes was prepared by Ficoll density gradient centrifugation (RD; I40650) and filtering through 35 µm nylon mesh. Cells were sequentially incubated with Zombie NIR (Biolegend 423106) in PBS (15 min at RT) for dead cell exclusion, rat anti-mouse CD16/32 (BD 553141, RRID:AB_394656, dilution 1:1000) in FACS buffer for Fc receptor blockade (10 min at 4°C), and fluorochrome-conjugated anti-mouse antibody mix diluted in Brilliant Stain Buffer (BD; 563794) for surface marker staining (30 min at 4°C). For intracellular staining, cells were fixed and permeabilized using the Foxp3 staining kit (ThermoFisher; 00-5523-00), followed by intracellular antibody incubation (O/N at 4°C). Data were acquired on Cytek Aurora (RRID:SCR_019826), BD FACSymphony A5 (RRID:SCR_022538), or BD FACS Canto II (RRID:SCR_018056) flow cytometer analyzers, and analyzed using FlowJo software (RRID:SCR_008520). The antibodies were diluted as follows: BUV615 anti-CD45 (BD 752418, RRID:AB_2917432, dilution 1:200), PerCP-Cy5.5 anti-B220 (BD 552771, RRID:AB_394457, dilution 1:200), PE-Cy7 anti-B220 (BD 552772, RRID:AB_394458, dilution 1:200), AF594 anti-B220 (Biolegend 103254, RRID:AB_2563229, dilution 1:200), BV786 anti-B220 (BD 563894, RRID:AB_2738472, dilution 1:200), BUV563 anti-CD38 (BD 741271, RRID:AB_2870812, dilution 1:250), BUV395 anti-CD38 (BD 740245, RRID:AB_2739992, dilution 1:250), APC-Cy7 anti-CD38 (Biolegend 102728, RRID:AB_2616968, dilution 1:200), BV421 anti-FAS/CD95 (BD 562633, RRID:AB_2737690, dilution 1:200), PE anti-FAS/CD95 (BD 554258, RRID:AB_395330, dilution 1:200), PE-Cy7 anti-FAS/CD95 (BD 557653, RRID:AB_396768, dilution 1:200), PE anti-CXCR4/CD184 (BD 561734, RRID:AB_11154227, dilution 1:125), PE-Cy7 anti-CD86 (BD 560582, RRID:AB_1727518, dilution 1:200), PerCP-Cy5.5 anti-CD45.1 (ThermoFisher 45-0453-82, dilution 1:200), AF647 anti-CD45.2 (Biolegend 109818, RRID:AB_492870, dilution 1:200), BUV737 anti-CD138 (BD 564430, RRID:AB_2738805, dilution 1:400), BV510 anti-IgD (BD 563110, RRID:AB_2737003, dilution 1:100), APC anti-CD4 (Biolegend 100412, RRID:AB_312697, dilution 1:200), PE anti-CD8 (Biolegend 100708, RRID:AB_312747, dilution 1:200).

To sort mouse GCB, CB and CC, GCB cells were first enriched from mouse splenocytes using the PNA MicroBead Kit (Miltenyi Biotec, 130-110-479), followed by antibody incubation and sorting on BD Aria II sorter (RRID:SCR_018091) with 5 lasers (355, 405, 488, 561, 640). DAPI was used to exclude dead cells.

### Immunohistochemistry, Immunofluorescence and Imaging of Tissues

Mouse organs (i.e., spleen, liver, lung, and kidney) were fixed in either 10% neutral buffered formalin for 24-48 hours at RT (for IHC) or 4% paraformaldehyde for 12 hours at 4°C (for IF), then transferred to 70% ethanol for storage before submitting to the core facility Laboratory of Comparative Pathology at Weill Cornell Medicine for paraffin embedding, sectioning and staining.

Briefly, five micron-sections cut using Microtome (Leica RM2255, RRID:SCR_020229) were deparaffinized and heat antigen retrieved in 10 mM sodium citrate buffer (pH 6.0) for 30 minutes using a steamer. For IHC, following antigen retrieval, endogenous peroxidase activity was blocked in 3% H2O2-methanol for 15 minutes at room temperature. Indirect IHC was performed using anti-species-specific biotinylated secondary antibodies followed by the introduction of avidin–horseradish peroxidase or avidin–alkaline phosphatase and developed by Vector Blue or DAB color substrates (Vector Laboratories). Sections were counterstained with hematoxylin. The following antibodies were used: Rat anti-mouse B220 (BD 550286, RRID:AB_393581), Rabbit anti-mouse CD4 (ab183685, RRID:AB_2686917), and Rabbit anti-mouse CD8 (ab217344, RRID:AB_2890649). For IF, the sections were stained with primary antibodies Rat Anti-Mouse B220 (BD 550286, RRID:AB_393581, 1:100 dilution) and Rabbit Anti-Mouse Ki67 (CST 12202, RRID:AB_2687446, 1:250 dilution) overnight at 4°C, followed by incubation with secondary antibodies Donkey anti-Rat IgG-AF488 (Invitrogen A21208, RRID:AB_141709, 1:500 dilution) and Donkey anti-Rabbit IgG-AF594 (Invitrogen A21207, RRID:AB_141637, 1:500 dilution) at room temperature for 1 hour. Autofluorescence in tissue sections due to aldehyde fixation was removed by treatment with the TrueVIEW Autofluorescence Quenching Kit for 4 min (Vector Laboratories; SP-8400-15). Sections were counterstained with DAPI (ThermoFisher 62247, 1:1000 dilution) for 5 min.

Slides were scanned using the All-in-One Fluorescence Microscope (KEYENCE, BZ-X810). Fiji software (ImageJ, RRID:SCR_003070) was used to quantify spleen and GC area size.

### Somatic Hypermutation Analysis of JH4 Intron

Genomic DNA of sorted GC B cells were isolated by Quick-DNA Microprep Kit (Zymo Research; D3020) according to the manufacturer’s protocol. JH4 intron sequences were amplified from genomic DNA by PCR with Phusion High-Fidelity DNA polymerase (NEB; M0530S) using JH4 forward primer and JH4 reverse primer. PCR products were resolved by agarose gel electrophoresis and JH4 intron sequences with size of 1.2 kb were extracted by QIAquick Gel Extraction Kit (Qiagen; 28706X4). DNA of JH4 sequences was ligated into the pCR-Blunt II-TOPO vector (ThermoFisher; K280002) and transformed into One Shot Chemically Competent *E. coli* according to the manufacturer’s instruction. Clones were sequenced by Sanger sequencing with JH4 sequencing primer. Primer sequences can be found in **Supplementary Table 6**.

### CRISPR/Cas9-mediated Gene Editing

To generate CREBBP-KO, CREBBP-R1446C, KMT2D-KO, and double loss-of-function isogenic OCI-Ly7 cells, ribonucleoprotein (RNP) complex containing Alt-R recombinant S.p. HiFi Cas9 Nuclease (IDT; 1081061), Alt-R CRISPR-Cas9 tracrRNA (IDT; 1072534), and Alt-R CRISPR-Cas9 crRNA targeting CREBBP exon 26 or KMT2D exon 4, were assembled in vitro following the manufacturer’s protocol and delivered into OCI-Ly7 cells by electroporation using SF Cell Line 96-well Nucleofector Kit (Lonza; V4SC-2096). ssODN for CREBBP R1446C mutagenesis was added along with RNP complex during electroporation as needed. Single cells were then seeded in 96-well plates by serial dilution. Clones were screened by Sanger sequencing of PCR amplicons and immunoblot. crRNA, ssODN and genotyping primer sequences can be found in **Supplementary Table 6**.

### Co-immunoprecipitation and Immunoblotting

Nuclear extracts were prepared as previously described (73). Nuclear extracts with salt adjustment (150 mM KCl) and detergent supplement (0.1% IGEPAL CA-630) were incubated with primary antibodies (CREBBP: Santa Cruz SC-369, RRID:AB_631006; KMT2D: MilliporeSigma ABE1867) overnight followed by addition of DynaBeads protein A (ThermoFisher; 10002D) the next day for 1.5 hours. After extensive washes for 6 times, the beads were boiled in 1X Laemmli sample buffer, separated by SDS-PAGE, and analyzed by immunoblot using the indicated primary antibodies.

For immunoblotting, cell lysates were resolved by SDS–PAGE, transferred to PVDF membrane, and probed with the following primary antibodies: anti-CREBBP (Santa Cruz SC-369, RRID:AB_631006, 1:1000) anti-KMT2D (MilliporeSigma ABE1867, 1:1000), anti-MED1 (ab64965, RRID:AB_1142301, 1:1000), anti-H3K4me1 (ab8895, RRID:AB_306847, 1:1000), anti-H3K27ac (ab4729, RRID:AB_2118291, 1:1000) and anti-H3 (ab18521, RRID:AB_732917, 1:1000). Membranes were then incubated with a peroxidase-conjugated correspondent secondary antibody and detected using enhanced chemiluminescence. Densitometry values were obtained using Fiji software (NIH).

### ELISA and ELISPOT

Mice were immunized with 100 ug NP19-OVA in alum via intraperitoneal injection. Serum collected at different days after immunization was subjected to ELISA. Briefly, serum was incubated in 96-well plates coated with NP28-BSA or NP8-BSA to capture low and high affinity NP-specific antibodies, respectively, followed by quantification of IgG1 abundance using the SBA Clonotyping System-HRP kit (SouthernBiotech; 5300-05) following the manufacturer’s instruction. For ELISPOT assay, bone marrow cells were collected on day 85 after immunization. 3 million BM cells were seeded into a 96-well MultiScreen IP Filter Plate (MilliporeSigma; MSIPS4510) coated with NP8-BSA and incubated overnight at 37°C. NP-specific LLPC spots were visualized using AP-conjugated goat anti-mouse IgG1 (SouthernBiotech; 1071-04) with enzyme substrates.

### Single-cell RNA-seq

Splenic B cells from mice immunized with SRBC for 10 days were pre-enriched by B220 MicroBeads (Miltenyi Biotec; 130-049-501) followed by flow sorting. Sorted B220+IgD-B cells from each spleen were submitted to the Epigenomics Core at Weill Cornell Medicine for library preparation and sequencing. Single-cell RNA-seq libraries were prepared using Chromium Single Cell 3’ Reagent Kit according to 10x Genomics specifications. Four independent single-cell suspensions (one per genotype) with a viability of 70-80% and a concentration of 400-800 cells/ul, were loaded onto the 10x Genomics Chromium platform to generate barcoded single-cell GEMs, targeting about 8000 single cells per sample. GEM-Reverse Transcription (53 °C for 45 min, 85 °C for 5 min; held at 4 °C) was performed in a C1000 Touch Thermal Cycler with 96-Deep Well Reaction Module (Bio-Rad). After RT reaction, GEMs were broken up and the single-strand cDNAs were cleaned up with DynaBeads MyOne Silane Beads (ThermoFisher; 37002D). The cDNAs were amplified for 12 cycles (98 °C for 3 min; 98 °C for 15 s, 63 °C for 20 s, 72 °C for 1 min). Quality of the cDNAs was assessed using Agilent Bioanalyzer 2100, obtaining an average product size of 1815 bp. These cDNAs were enzymatically fragmented, end repaired, A-tailed, subjected to a double-sided size selection with SPRIselect beads (Beckman Coulter) and ligated to adaptors provided in the kit. A unique sample index for each library was introduced through 14 cycles of PCR amplification using the indexes provided in the kit (98 °C for 45 s; 98 °C for 20 s, 54 °C for 30 s, and 72 °C for 20 s x 14 cycles; 72 °C for 1 min; held at 4 °C). Indexed libraries were subjected to a second double-sided size selection, and libraries were then quantified using Qubit (ThermoFisher, RRID:SCR_020553). The quality was assessed on an Agilent Bioanalyzer 2100 (RRID:SCR_019389), obtaining an average library size of 455 bp.

Libraries were diluted to 2 nM and clustered on an Illumina NovaSeq 6000 (RRID:SCR_016387) on a pair-end read flow cell and sequenced for 28 cycles on R1 (10x barcode and the UMIs), followed by 8 cycles of I7 Index (sample Index), and 98 bases on R2 (transcript), with a coverage around 200M reads per sample. Primary processing of sequencing images was done using Illumina’s Real Time Analysis software (RTA).

### Single-cell RNA-seq Analysis

Fastq files were processed with cellranger version 3.0.0. This data was combined with previously published single cell datasets (H1, Ezh2 and Smc3) and processed using Seurat version 4.0 (74). Cell types of these integrated datasets were assigned together from previous annotations using the Seurat TransferLabel function and previous reference datasets (37). Proper assignment of clusters was assessed by both individual genes, and previously published gene signatures. Gene signature module scores were calculated using the AddModuleScore function, with a control value of 5. Cells from this experiment were then subsetted and reprocessed using 5000 variable genes to generate the UMAP with the appropriate cell populations. Cell type percentages were calculated after removal of non-GC cells and significance was tested using a fisher’s exact test. Pseudotime trajectories were generated with slingshot 2.4.0 (38) on GC B cells. Density of cells along this pseudotime was plotted, and significance was tested using wilcoxon rank sum. Gene signatures were plotted against pseudotime and broken up into 10 deciles. Expression of the gene signatures in each of these deciles were tested using wilcoxon rank sum between the WT and mutant cells. The differences in the splines were plotted and colored according to decile significance.

### Bulk RNA-seq

Total RNAs were extracted from sorted cell suspensions using TRIzol LS Reagent (Invitrogen,10296010) following the manufacturer’s instructions. RNA concentration was determined using the Qubit RNA High Sensitivity Kit (ThermoFisher, Q32855) and integrity was assessed using RNA 6000 Pico Kit (Agilent, 5067-1513) run on the Agilent 2100 Bioanalyzer. 100ng-400ng samples with RNA Integrity Number (RIN) > 9 were submitted to MedGenome for library preparation using the Illumina TruSeq Stranded mRNA Library Kit and PE100 sequencing on NovaSeq.

### Bulk RNA-seq Analysis

Fastq files were processed with the nfcore RNA-seq pipeline version 2.0, aligned to mm10 or hg38. Count files from the nfcore pipeline output were processed with DESeq2. PCA clustering was performed on VST expression values for all GENCODE vM25 genes. Hierarchical clustering was then performed on the top 10% most variable genes using Euclidean distance and Ward’s method. DESeq2 was also used to determine DEGs. Transcripts with less than 10 combined reads among all replicates or that DESeq2 failed to generate a p value for were removed from the datasets. *P* values were readjusted using Benjamini-Hochberg correction and differentially expressed genes were defined using a fold change < 1.5 fcutoff and padj < 0.01. Enrichment of signatures was performed using iPAGE. Briefly, unsupervised pathway analysis was performed using information-theoretic pathway analysis approach as described in (75). Briefly, pathways that are informative about non-overlapping gene groups were identified. Pathways annotations were used from the Biological Process annotations of the Gene Ontology database (http://www.geneontology.org) and signature categories from the Staudt Lab Signature database (76). Only human-curated annotations were used from the Gene Ontology database and only pathways with 5 genes or more, and with 300 genes or less were evaluated. This pathway analysis estimates how informative each pathway is about the target gene groups, and applies a randomization-based statistical test to assess the significance of the highest information values. We use the default significance threshold of p<0.005. We estimated the false discovery rate (FDR) by randomizing the input profiles iteratively on shuffled profiles with identical parameters and thresholds, finding that the FDR was always less than 5%. For each informative pathway, we determined the extent to which the pathway was over-represented in the target gene group, using the hypergeometric distribution, as described in (77). Clustering to define modules was performed on standardized log2 fold-change values relative to WT CB cells using fuzzy c-means clustering with 8 clusters and fuzzifier parameter as selected by Schwämmle-Jensen method (78).

### Bulk ATAC-seq

Bulk ATAC-seq libraries were prepared from sorted cell suspensions following the Omni-ATAC protocol (79). Briefly, 60,000 freshly sorted live cells were washed in cold PBS, lysed in 60 ul cold lysis buffer (10 mM Tris-HCl, pH 7.5, 10 mM NaCl, 3 mM MgCl2, 0.1% NP-40, 0.1% Tween-20, and 0.01% Digitonin) for 3 min on ice, followed by addition of 1 ml wash buffer (10 mM Tris-HCl, pH 7.5, 10 mM NaCl, 3 mM MgCl2, and 0.1% Tween-20) and centrifugation to pellet nuclei. Nuclei were resuspended in a transposition reaction mix prepared using the Illumina Tagment DNA Enzyme and Buffer Kit (Illumina, 20034197) and incubated at 37 °C for 30 minutes on thermomixer at 1,000 rpm to allow for transposition-based DNA fragmentation and adapter ligation. Tagged DNA fragments were purified using the Clean & Concentrator-5 Kit (ZYMO, D4014), then subjected to PCR amplification and double-sided bead purification (to remove primer dimers and larger than 1,000 bp fragments) using the AMPure XP beads (Beckman Coulter, A63881). Library size distribution and quality was measured using the High Sensitivity DNA Kit (Agilent, 5067-4626) run on the Agilent 2100 Bioanalyzer. Libraries were submitted to MedGenome for PE50 sequencing on NovaSeq.

### Bulk ATAC-seq Analysis

Fastq files were processed with the nfcore ATAC-seq pipeline version 1.2.1 and aligned to mm10. Count files were processed the same way as the RNA-seq data to generate PCA plots and hierarchical clustering, with male/female included as a covariate. Differential peaks were also defined the same way as the RNA-seq, with the Benjamini-Hochberg adjusted pvalue < 0.01 and with a log2(1.5) fold change cutoff. These differential peaks were grouped according to k-means clustering, using clustGap function of the cluster R package (v2.1.0; Maechler, M., Rousseeuw, P., Struyf, A., Hubert, M., & Hornik, K. (2019). cluster: Cluster Analysis Basics and Extensions.) to calculate the optimal number of clusters. Supervised clustering was then performed by merging clusters of similar patterning across three genotypes. A fisher’s exact test was then used to determine the odds ratio of the genomic feature of these differential peaks, using distance to TSS (+/-5kb) to define promoters, and H3K27ac or H3K4me1 to define putative poised or active enhancer types. Super-enhancers were defined using the ROSE algorithm (72, 80) on previously published H3K27ac Mint-ChIP data, excluding any peaks within 2500bp of a protein coding RNA transcription start site (TSS).

For RNAseq GSEA, gene expression was linked to differential ATACseq peaks using GREAT regulatory regions by basal plus extension model (48). For ATACseq GSEA, peaks were linked to differentially expressed genes by previously defined mm10 germinal center TADs (H1). Constituent peaks of super-enhancers were used to determine differential accessibility at enhancers or super-enhancers, and significance was tested with wilcoxon rank sum. For the ranked Super-Enhancer accessibility, constituent peaks (overlap with the H3K27Ac super-enhancer peaks of at least 1 base pair) were summed per super-enhancer, then DESeq2 was used to normalize these counts with the size factors from the total ATACseq to account for read depth.

Peak clusters were determined by taking the variance stabilized transformed (VST) normalized counts in these merged regions for Mint-ChIP signal in H3K27Ac, H3K4me1, H3K4me3, and the ATAC-seq data from the four genotypes. t-SNE plots of these data were generated, and k-means clustering was used to define clusters. The clusters were linked with genes using distance to the closest TSS. The distance to TSS and the expression of these genes were used as a read-out of the cluster results.

TF motif enrichment was calculated as previously described (50). Mainly, JASPAR mammalian motifs were scanned for in the ATACseq peaks, and the log2 fold-change of the peak accessibility change was linearly modeled as log2FC ∼ β_0_ + x_1_β_0_ + ⋯ + x_n_β_n_ + GC ∗ β_GC_, where x*_i_* denotes a boolean whether the i^th^ transcription factor is present in the peak or not, and GC is the GC content of the peak. The effect sizes and p-values for each term are taken after multivariate linear modeling using ordinary least squares regression with robust standard errors.

### CUT&RUN

OCI-Ly7 cells were subjected to CUT&RUN assay according to the EpiCypher CUTANA CUT&RUN Protocol (81). In brief, 500K cells were immobilized onto activated ConA magnetic beads (Bangs Laboratories, BP531), followed by permeabilization and incubation with 0.5 ug of antibodies (H3K4me1, 13-0040, RRID:AB_2923143; H3K4me3, 13-0028; or H3K27ac, 13-0045, RRID:AB_2923489; SNAP-Certified for CUT&RUN and ChIP by EpiCypher) overnight at 4 °C. On the next day, the cell-bead slurry was washed and incubated with pAG-MNase (1:20 dilution, EpiCypher, 15-1116) for 10 min at RT. MNase was then activated by addition of CaCl2 to cleave targeted chromatin for 2 h at 4 °C. After chromatin digestion, MNase activity was stopped and chromatin fragments released into supernatant were purified using the NEB Monarch DNA Cleanup Kit (NEB, T1030) per manufacturer’s instruction. 10 ng DNA was subjected to library preparation using NEBNext Ultra II DNA Library Prep Kit for Illumina (NEB, E7645) according to the manufacturer’s protocol. Libraries were loaded onto the Illumina HiSeq for pair-end 150 bp sequencing with a sequencing depth of around 6M reads per sample.

### Mint-ChIP sequencing

Mint-ChIP was performed using anti-H3K4me1 (Abcam ab8895, RRID:AB_306847) and anti-H3K4me3 (Abcam ab8580, RRID:AB_306649) as previously described (82). Briefly, 5 × 10^4^ flow-sorted mouse GC B cells in triplicate were processed with MNase to fragment native chromatin. Barcoded adaptors were ligated to every sample, and samples were multiplexed for ChIP. After ChIP, material was linearly amplified by RNA in vitro transcription in the presence of the RNase inhibitor RNaseOUT (Invitrogen, 10777019). Fragments were reverse transcribed and amplified as a library. Sequencing was run on Illumina NextSeq 500.

### ChIP-seq and CUT&RUN Analysis

Previously published CREBBP and KMT2D ChIP-seq data was aligned to hg38, with peaks being called with MACS2. The number of overlapped ChIP-seq peaks were calculated using the bedtools intersect function. Cut&Run fastq files were processed with nfcore cutandrun version 2.0, with IGG controls used per genotype. Read depth was normalized to reads at promoters. Peaks were called with SEACR and summits were called with MACS2. Counts were processed with DiffBind 3.6.1 (83). The union of consensus peaks from all marks was taken. Reads under the merged consensus peak was calculated for each mark by SubRead featureCounts (84) and normalized by variance stabilized transform (VST) (85). t-SNE was generated using the Cut&Run signal from all histone marks for baseline WT samples and additional features of log2 fold-changes of each genotype compared to WT. The regions were clustered using k-means clustering. RNA expression was linked by taking the closest gene transcription start site (TSS) to each peak. Super-enhancers were defined using ROSE on the WT H3K27Ac Cut&Run data, with non-coding RNA TSS removed and default settings (2500bp from TSS). Cut&Run heatmaps were generated with deepTools 3.5.1 (86). For H3K4me1, peak summits that overlapped with Cluster 1 peaks were used to generate peaks. For H3K27ac, peak center of peaks called by SEACR were used. RNA expression for both Cut&Run and CHIP was linked by GREAT and previously defined TADs.

### ChIP-qPCR

CREBBP and KMT2D ChIP assays were performed as previously described (19, 20). 50 million cells were first cross-linked with 2 mM DSG (disuccinimidyl glutarate, ThermoFisher, 20593) for 45 min at RT, and then 1% formaldehyde for 10 min at RT in PBS. Fixed cells were lysed to isolate nuclei, followed by sonication, incubation with Dynabeads (ThermoFisher, 11204D) coated with CREBBP (Santa Cruz, SC-369, RRID:AB_631006) or KMT2D antibody (Millipore, ABE1867), wash, elution, reverse-crosslink, and DNA purification. The recovered ChIP and input DNA was amplified by real-time quantitative PCR using SYBR Green (Applied Biosystems, 4385614) on QuantStudio 6 Flex Real-Time PCR System (Applied Biosystems). Data was analyzed by percent input method. ChIP-qPCR primers were listed in **Supplementary Table 6**.

### RT-qPCR

cDNA was prepared from extracted RNA using cDNA synthesis kit (ThermoFisher Scientific, K1641) and detected by fast SYBR Green (Applied Biosystems, 4385614) on QuantStudio 6 Flex Real-Time PCR System (Applied Biosystems) using the primers listed in **Supplementary Table 6**. qPCR signal for each gene was normalized to those of Gapdh (mouse) or HPRT (human) using the ΔCT method. Results were represented as fold expression relative to WT with the standard deviation for 3-4 biological replicates.

## Statistical Analysis

Statistical methods used for p value calculation were specified in figure legends for the corresponding panels. Statistical analysis was conducted using GraphPad Prism 9 (RRID:SCR_002798) or the R statistical language scripts (RRID:SCR_001905) and packages specified. Data were judged to be statistically significant when p < 0.05.

## Data Availability

All genomics data that support the findings of this study have been deposited in the NCBI Gene Expression Omnibus (GEO) with accession number GSE224513. All other data are available from the corresponding authors upon request.

## Author’s Contributions

**J.Li:** Conceptualization, resources, data curation, formal analysis, validation, investigation, visualization, methodology, writing–original draft, writing–review and editing. **C.R.Chin:** Conceptualization, resources, data curation, formal analysis, validation, investigation, visualization, methodology, writing–original draft, writing–review and editing. **H.-Y.Ying**: Conceptualization, resources, data curation, formal analysis, validation, investigation, visualization, methodology, writing–review and editing. **C.Meydan:** data curation, formal analysis, visualization, methodology, writing–review and editing. **M.R.Teater:** data curation, formal analysis, visualization, methodology, writing– review and editing. **M.Xia**: investigation, writing–review and editing. **P.Farinha:** investigation, visualization, methodology, writing–review and editing. **K.Takata:** investigation, visualization, methodology, writing–review and editing. **C.-S.Chu:** investigation, writing–review and editing. **M.A.Rivas:** investigation, writing–review and editing. **A.Chadburn:** investigation, visualization, writing–review and editing. **C.Steidl:** resources, writing–review and editing. **D.W.Scott:** resources, writing–review and editing. **R.G.Roeder:** resources, writing–review and editing. **C.E.Mason:** resources, supervision, funding acquisition, writing–review and editing. **W.Béguelin:** Conceptualization, resources, formal analysis, supervision, funding acquisition, methodology, writing– original draft, project administration, writing–review and editing. **A.M.Melnick:** Conceptualization, resources, formal analysis, supervision, funding acquisition, methodology, writing–original draft, project administration, writing–review and editing.

## Acknowledgements

We thank Dr. Kai Ge (NIH) for providing the *Kmt2d* floxed mice used in this study. We especially thank Dr. Martin T. Wells (Cornell University) for advice on statistical analysis. We thank all the members in the Melnick lab for their helpful discussion and technical support. In addition, we thank Weill Cornell Medicine core facilities, including Epigenomics Core, Genomics Resources Core, Flow Cytometry Core, and Laboratory of Comparative Pathology for their professional support of this work. This work is supported by grants A.M.M.: R35 CA220499, LLS SCOR 7021-20, IFLI collaborative research award. W.B. is supported by NCI R01 CA270245, ASH Junior Faculty Scholar Award, Leukemia & Lymphoma Society, Lymphoma Research Foundation, and The Follicular Lymphoma Foundation. C.R.C. is supported by 5 F31 CA254302-02. H.-Y.Y. was supported by AACR-Takeda Oncology Fellowship in Lymphoma Research. M.A.R. is supported by the ASH Junior Faculty Scholar Award. This work is supported by program project grant funding from the Terry Fox Research Institute (grant 1061) and the BC Cancer Foundation. D.W.S. is supported by a Michael Smith Foundation for Health Research Health Professional Investigator award (18646). R.G.R. was supported by grants R01AI148387 and LLS SCOR 7021-20. C.E.M. thanks the Scientific Computing Unit (SCU) at WCM, the WorldQuant Foundation, the National Institutes of Health (R01MH117406, R01CA249054, P01CA214274), and the LLS (MCL7001-18, LLS-9238-16, LLS-MCL7001-18, LLS 7029-23).

## REFERENCES

1. Baylin SB, Jones PA. Epigenetic Determinants of Cancer. Cold Spring Harb Perspect Biol 2016;8(9) doi 10.1101/cshperspect.a019505.

2. Dawson MA, Kouzarides T. Cancer epigenetics: from mechanism to therapy. Cell 2012;150(1):12–27 doi 10.1016/j.cell.2012.06.013.

3. Duy C, Beguelin W, Melnick A. Epigenetic Mechanisms in Leukemias and Lymphomas. Cold Spring Harb Perspect Med 2020;10(12) doi 10.1101/cshperspect.a034959.

4. Allis CD, Jenuwein T. The molecular hallmarks of epigenetic control. Nat Rev Genet 2016;17(8):487–500 doi 10.1038/nrg.2016.59.

5. Okosun J, Bodor C, Wang J, Araf S, Yang CY, Pan C, et al. Integrated genomic analysis identifies recurrent mutations and evolution patterns driving the initiation and progression of follicular lymphoma. Nat Genet 2014;46(2):176–81 doi 10.1038/ng.2856.

6. Pasqualucci L, Khiabanian H, Fangazio M, Vasishtha M, Messina M, Holmes AB, et al. Genetics of follicular lymphoma transformation. Cell Rep 2014;6(1):130–40 doi 10.1016/j.celrep.2013.12.027.

7. Chapuy B, Stewart C, Dunford AJ, Kim J, Kamburov A, Redd RA, et al. Molecular subtypes of diffuse large B cell lymphoma are associated with distinct pathogenic mechanisms and outcomes. Nat Med 2018;24(5):679–90 doi 10.1038/s41591-018-0016-8.

8. Mlynarczyk C, Fontan L, Melnick A. Germinal center-derived lymphomas: The darkest side of humoral immunity. Immunol Rev 2019;288(1):214–39 doi 10.1111/imr.12755.

9. Schmitz R, Wright GW, Huang DW, Johnson CA, Phelan JD, Wang JQ, et al. Genetics and Pathogenesis of Diffuse Large B-Cell Lymphoma. N Engl J Med 2018;378(15):1396–407 doi 10.1056/NEJMoa1801445.

10. Green MR, Gentles AJ, Nair RV, Irish JM, Kihira S, Liu CL, et al. Hierarchy in somatic mutations arising during genomic evolution and progression of follicular lymphoma. Blood 2013;121(9):1604–11 doi 10.1182/blood-2012-09-457283.

11. Green MR, Kihira S, Liu CL, Nair RV, Salari R, Gentles AJ, et al. Mutations in early follicular lymphoma progenitors are associated with suppressed antigen presentation. Proc Natl Acad Sci U S A 2015;112(10):E1116–25 doi 10.1073/pnas.1501199112.

12. Calo E, Wysocka J. Modification of enhancer chromatin: what, how, and why? Mol Cell 2013;49(5):825–37 doi 10.1016/j.molcel.2013.01.038.

13. Froimchuk E, Jang Y, Ge K. Histone H3 lysine 4 methyltransferase KMT2D. Gene 2017;627:337–42 doi 10.1016/j.gene.2017.06.056.

14. Tie F, Banerjee R, Stratton CA, Prasad-Sinha J, Stepanik V, Zlobin A, et al. CBP-mediated acetylation of histone H3 lysine 27 antagonizes Drosophila Polycomb silencing. Development 2009;136(18):3131–41 doi 10.1242/dev.037127.

15. Creyghton MP, Cheng AW, Welstead GG, Kooistra T, Carey BW, Steine EJ, et al. Histone H3K27ac separates active from poised enhancers and predicts developmental state. Proc Natl Acad Sci U S A 2010;107(50):21931–6 doi 10.1073/pnas.1016071107.

16. Raisner R, Kharbanda S, Jin L, Jeng E, Chan E, Merchant M, et al. Enhancer Activity Requires CBP/P300 Bromodomain-Dependent Histone H3K27 Acetylation. Cell Rep 2018;24(7):1722–9 doi 10.1016/j.celrep.2018.07.041.

17. Garcia-Ramirez I, Tadros S, Gonzalez-Herrero I, Martin-Lorenzo A, Rodriguez-Hernandez G, Moore D, et al. Crebbp loss cooperates with Bcl2 overexpression to promote lymphoma in mice. Blood 2017;129(19):2645–56 doi 10.1182/blood-2016-08-733469.

18. Hashwah H, Schmid CA, Kasser S, Bertram K, Stelling A, Manz MG, et al. Inactivation of CREBBP expands the germinal center B cell compartment, down-regulates MHCII expression and promotes DLBCL growth. Proc Natl Acad Sci U S A 2017;114(36):9701–6 doi 10.1073/pnas.1619555114.

19. Jiang Y, Ortega-Molina A, Geng H, Ying HY, Hatzi K, Parsa S, et al. CREBBP Inactivation Promotes the Development of HDAC3-Dependent Lymphomas. Cancer Discov 2017;7(1):38–53 doi 10.1158/2159-8290.CD-16-0975.

20. Ortega-Molina A, Boss IW, Canela A, Pan H, Jiang Y, Zhao C, et al. The histone lysine methyltransferase KMT2D sustains a gene expression program that represses B cell lymphoma development. Nat Med 2015;21(10):1199–208 doi 10.1038/nm.3943.

21. Zhang J, Dominguez-Sola D, Hussein S, Lee JE, Holmes AB, Bansal M, et al. Disruption of KMT2D perturbs germinal center B cell development and promotes lymphomagenesis. Nat Med 2015;21(10):1190–8 doi 10.1038/nm.3940.

22. Zhang J, Vlasevska S, Wells VA, Nataraj S, Holmes AB, Duval R, et al. The CREBBP Acetyltransferase Is a Haploinsufficient Tumor Suppressor in B-cell Lymphoma. Cancer Discov 2017;7(3):322–37 doi 10.1158/2159-8290.CD-16-1417.

23. Mesin L, Ersching J, Victora GD. Germinal Center B Cell Dynamics. Immunity 2016;45(3):471–82 doi 10.1016/j.immuni.2016.09.001.

24. Young C, Brink R. The unique biology of germinal center B cells. Immunity 2021;54(8):1652–64 doi 10.1016/j.immuni.2021.07.015.

25. Figueroa ME, Abdel-Wahab O, Lu C, Ward PS, Patel J, Shih A, et al. Leukemic IDH1 and IDH2 mutations result in a hypermethylation phenotype, disrupt TET2 function, and impair hematopoietic differentiation. Cancer Cell 2010;18(6):553–67 doi 10.1016/j.ccr.2010.11.015.

26. Ma MCJ, Tadros S, Bouska A, Heavican T, Yang H, Deng Q, et al. Subtype-specific and co-occurring genetic alterations in B-cell non-Hodgkin lymphoma. Haematologica 2022;107(3):690–701 doi 10.3324/haematol.2020.274258.

27. Shih AH, Jiang Y, Meydan C, Shank K, Pandey S, Barreyro L, et al. Mutational cooperativity linked to combinatorial epigenetic gain of function in acute myeloid leukemia. Cancer Cell 2015;27(4):502–15 doi 10.1016/j.ccell.2015.03.009.

28. Lacy SE, Barrans SL, Beer PA, Painter D, Smith AG, Roman E, et al. Targeted sequencing in DLBCL, molecular subtypes, and outcomes: a Haematological Malignancy Research Network report. Blood 2020;135(20):1759–71 doi 10.1182/blood.2019003535.

29. Wright GW, Huang DW, Phelan JD, Coulibaly ZA, Roulland S, Young RM, et al. A Probabilistic Classification Tool for Genetic Subtypes of Diffuse Large B Cell Lymphoma with Therapeutic Implications. Cancer Cell 2020;37(4):551–68 e14 doi 10.1016/j.ccell.2020.03.015.

30. Han G, Deng Q, Marques-Piubelli ML, Dai E, Dang M, Ma MCJ, et al. Follicular Lymphoma Microenvironment Characteristics Associated with Tumor Cell Mutations and MHC Class II Expression. Blood Cancer Discov 2022;3(5):428–43 doi 10.1158/2643-3230.BCD-21-0075.

31. Casola S, Cattoretti G, Uyttersprot N, Koralov SB, Seagal J, Hao Z, et al. Tracking germinal center B cells expressing germ-line immunoglobulin gamma1 transcripts by conditional gene targeting. Proc Natl Acad Sci U S A 2006;103(19):7396–401 doi 10.1073/pnas.0602353103.

32. Kang-Decker N, Tong C, Boussouar F, Baker DJ, Xu W, Leontovich AA, et al. Loss of CBP causes T cell lymphomagenesis in synergy with p27Kip1 insufficiency. Cancer Cell 2004;5(2):177–89 doi 10.1016/s1535-6108(04)00022-4.

33. Lee JE, Wang C, Xu S, Cho YW, Wang L, Feng X, et al. H3K4 mono- and di-methyltransferase MLL4 is required for enhancer activation during cell differentiation. Elife 2013;2:e01503 doi 10.7554/eLife.01503.

34. Egle A, Harris AW, Bath ML, O’Reilly L, Cory S. VavP-Bcl2 transgenic mice develop follicular lymphoma preceded by germinal center hyperplasia. Blood 2004;103(6):2276–83 doi 10.1182/blood-2003-07-2469.

35. Ogilvy S, Metcalf D, Print CG, Bath ML, Harris AW, Adams JM. Constitutive Bcl-2 expression throughout the hematopoietic compartment affects multiple lineages and enhances progenitor cell survival. Proc Natl Acad Sci U S A 1999;96(26):14943–8 doi 10.1073/pnas.96.26.14943.

36. Boothby M, Rickert RC. Metabolic Regulation of the Immune Humoral Response. Immunity 2017;46(5):743–55 doi 10.1016/j.immuni.2017.04.009.

37. Rivas MA, Durmaz C, Kloetgen A, Chin CR, Chen Z, Bhinder B, et al. Cohesin Core Complex Gene Dosage Contributes to Germinal Center Derived Lymphoma Phenotypes and Outcomes. Front Immunol 2021;12:688493 doi 10.3389/fimmu.2021.688493.

38. Street K, Risso D, Fletcher RB, Das D, Ngai J, Yosef N, et al. Slingshot: cell lineage and pseudotime inference for single-cell transcriptomics. BMC Genomics 2018;19(1):477 doi 10.1186/s12864-018-4772-0.

39. Furlong EEM, Levine M. Developmental enhancers and chromosome topology. Science 2018;361(6409):1341–5 doi 10.1126/science.aau0320.

40. Rivas MA, Meydan C, Chin CR, Challman MF, Kim D, Bhinder B, et al. Smc3 dosage regulates B cell transit through germinal centers and restricts their malignant transformation. Nat Immunol 2021;22(2):240–53 doi 10.1038/s41590-020-00827-8.

41. Laidlaw BJ, Cyster JG. Transcriptional regulation of memory B cell differentiation. Nat Rev Immunol 2021;21(4):209–20 doi 10.1038/s41577-020-00446-2.

42. Song S, Matthias PD. The Transcriptional Regulation of Germinal Center Formation. Front Immunol 2018;9:2026 doi 10.3389/fimmu.2018.02026.

43. Arenzana TL, Smith-Raska MR, Reizis B. Transcription factor Zfx controls BCR-induced proliferation and survival of B lymphocytes. Blood 2009;113(23):5857–67 doi 10.1182/blood-2008-11-188888.

44. Banerjee A, Sindhava V, Vuyyuru R, Jha V, Hodewadekar S, Manser T, et al. YY1 Is Required for Germinal Center B Cell Development. PLoS One 2016;11(5):e0155311 doi 10.1371/journal.pone.0155311.

45. Chen J, Bardes EE, Aronow BJ, Jegga AG. ToppGene Suite for gene list enrichment analysis and candidate gene prioritization. Nucleic Acids Res 2009;37(Web Server issue):W305–11 doi 10.1093/nar/gkp427.

46. Kuleshov MV, Jones MR, Rouillard AD, Fernandez NF, Duan Q, Wang Z, et al. Enrichr: a comprehensive gene set enrichment analysis web server 2016 update. Nucleic Acids Res 2016;44(W1):W90–7 doi 10.1093/nar/gkw377.

47. Hnisz D, Abraham BJ, Lee TI, Lau A, Saint-Andre V, Sigova AA, et al. Super-enhancers in the control of cell identity and disease. Cell 2013;155(4):934–47 doi 10.1016/j.cell.2013.09.053.

48. McLean CY, Bristor D, Hiller M, Clarke SL, Schaar BT, Lowe CB, et al. GREAT improves functional interpretation of cis-regulatory regions. Nat Biotechnol 2010;28(5):495–501 doi 10.1038/nbt.1630.

49. Bunting KL, Soong TD, Singh R, Jiang Y, Beguelin W, Poloway DW, et al. Multi-tiered Reorganization of the Genome during B Cell Affinity Maturation Anchored by a Germinal Center-Specific Locus Control Region. Immunity 2016;45(3):497–512 doi 10.1016/j.immuni.2016.08.012.

50. Doane AS, Chu CS, Di Giammartino DC, Rivas MA, Hellmuth JC, Jiang Y, et al. OCT2 pre-positioning facilitates cell fate transition and chromatin architecture changes in humoral immunity. Nat Immunol 2021;22(10):1327–40 doi 10.1038/s41590-021-01025-w.

51. Kridel R, Chan FC, Mottok A, Boyle M, Farinha P, Tan K, et al. Histological Transformation and Progression in Follicular Lymphoma: A Clonal Evolution Study. PLoS Med 2016;13(12):e1002197 doi 10.1371/journal.pmed.1002197.

52. Mondello P, Tadros S, Teater M, Fontan L, Chang AY, Jain N, et al. Selective Inhibition of HDAC3 Targets Synthetic Vulnerabilities and Activates Immune Surveillance in Lymphoma. Cancer Discov 2020;10(3):440–59 doi 10.1158/2159-8290.CD-19-0116.

53. Hatzi K, Jiang Y, Huang C, Garrett-Bakelman F, Gearhart MD, Giannopoulou EG, et al. A hybrid mechanism of action for BCL6 in B cells defined by formation of functionally distinct complexes at enhancers and promoters. Cell Rep 2013;4(3):578–88 doi 10.1016/j.celrep.2013.06.016.

54. Huang C, Hatzi K, Melnick A. Lineage-specific functions of Bcl-6 in immunity and inflammation are mediated by distinct biochemical mechanisms. Nat Immunol 2013;14(4):380–8 doi 10.1038/ni.2543.

55. Ranuncolo SM, Polo JM, Dierov J, Singer M, Kuo T, Greally J, et al. Bcl-6 mediates the germinal center B cell phenotype and lymphomagenesis through transcriptional repression of the DNA-damage sensor ATR. Nat Immunol 2007;8(7):705–14 doi 10.1038/ni1478.

56. Polo JM, Ci W, Licht JD, Melnick A. Reversible disruption of BCL6 repression complexes by CD40 signaling in normal and malignant B cells. Blood 2008;112(3):644–51 doi 10.1182/blood-2008-01-131813.

57. Wang SP, Tang Z, Chen CW, Shimada M, Koche RP, Wang LH, et al. A UTX-MLL4-p300 Transcriptional Regulatory Network Coordinately Shapes Active Enhancer Landscapes for Eliciting Transcription. Mol Cell 2017;67(2):308–21 e6 doi 10.1016/j.molcel.2017.06.028.

58. Pradeepa MM. Causal role of histone acetylations in enhancer function. Transcription 2017;8(1):40–7 doi 10.1080/21541264.2016.1253529.

59. Ooi SK, Qiu C, Bernstein E, Li K, Jia D, Yang Z, et al. DNMT3L connects unmethylated lysine 4 of histone H3 to de novo methylation of DNA. Nature 2007;448(7154):714–7 doi 10.1038/nature05987.

60. Local A, Huang H, Albuquerque CP, Singh N, Lee AY, Wang W, et al. Identification of H3K4me1-associated proteins at mammalian enhancers. Nat Genet 2018;50(1):73–82 doi 10.1038/s41588-017-0015-6.

61. Yan J, Chen SA, Local A, Liu T, Qiu Y, Dorighi KM, et al. Histone H3 lysine 4 monomethylation modulates long-range chromatin interactions at enhancers. Cell Res 2018;28(2):204–20 doi 10.1038/cr.2018.1.

62. Park YK, Lee JE, Yan Z, McKernan K, O’Haren T, Wang W, et al. Interplay of BAF and MLL4 promotes cell type-specific enhancer activation. Nat Commun 2021;12(1):1630 doi 10.1038/s41467-021-21893-y.

63. Fasciani A, D’Annunzio S, Poli V, Fagnocchi L, Beyes S, Michelatti D, et al. MLL4-associated condensates counterbalance Polycomb-mediated nuclear mechanical stress in Kabuki syndrome. Nat Genet 2020;52(12):1397–411 doi 10.1038/s41588-020-00724-8.

64. Sabari BR, Dall’Agnese A, Boija A, Klein IA, Coffey EL, Shrinivas K, et al. Coactivator condensation at super-enhancers links phase separation and gene control. Science 2018;361(6400) doi 10.1126/science.aar3958.

65. Weinert BT, Narita T, Satpathy S, Srinivasan B, Hansen BK, Scholz C, et al. Time-Resolved Analysis Reveals Rapid Dynamics and Broad Scope of the CBP/p300 Acetylome. Cell 2018;174(1):231–44 e12 doi 10.1016/j.cell.2018.04.033.

66. Beguelin W, Teater M, Gearhart MD, Calvo Fernandez MT, Goldstein RL, Cardenas MG, et al. EZH2 and BCL6 Cooperate to Assemble CBX8-BCOR Complex to Repress Bivalent Promoters, Mediate Germinal Center Formation and Lymphomagenesis. Cancer Cell 2016;30(2):197–213 doi 10.1016/j.ccell.2016.07.006.

67. Esensten JH, Helou YA, Chopra G, Weiss A, Bluestone JA. CD28 Costimulation: From Mechanism to Therapy. Immunity 2016;44(5):973–88 doi 10.1016/j.immuni.2016.04.020.

68. Cannons JL, Qi H, Lu KT, Dutta M, Gomez-Rodriguez J, Cheng J, et al. Optimal germinal center responses require a multistage T cell:B cell adhesion process involving integrins, SLAM-associated protein, and CD84. Immunity 2010;32(2):253–65 doi 10.1016/j.immuni.2010.01.010.

69. Liu D, Xu H, Shih C, Wan Z, Ma X, Ma W, et al. T-B-cell entanglement and ICOSL-driven feed-forward regulation of germinal centre reaction. Nature 2015;517(7533):214–8 doi 10.1038/nature13803.

70. Reimer D, Lee AY, Bannan J, Fromm P, Kara EE, Comerford I, et al. Early CCR6 expression on B cells modulates germinal centre kinetics and efficient antibody responses. Immunol Cell Biol 2017;95(1):33–41 doi 10.1038/icb.2016.68.

71. Heward J, Konali L, D’Avola A, Close K, Yeomans A, Philpott M, et al. KDM5 inhibition offers a novel therapeutic strategy for the treatment of KMT2D mutant lymphomas. Blood 2021;138(5):370–81 doi 10.1182/blood.2020008743.

72. Whyte WA, Orlando DA, Hnisz D, Abraham BJ, Lin CY, Kagey MH, et al. Master transcription factors and mediator establish super-enhancers at key cell identity genes. Cell 2013;153(2):307–19 doi 10.1016/j.cell.2013.03.035.

73. Folco EG, Lei H, Hsu JL, Reed R. Small-scale nuclear extracts for functional assays of gene-expression machineries. J Vis Exp 2012(64) doi 10.3791/4140.

74. Hao Y, Hao S, Andersen-Nissen E, Mauck WM, 3rd, Zheng S, Butler A, et al. Integrated analysis of multimodal single-cell data. Cell 2021;184(13):3573–87 e29 doi 10.1016/j.cell.2021.04.048.

75. Goodarzi H, Elemento O, Tavazoie S. Revealing global regulatory perturbations across human cancers. Mol Cell 2009;36(5):900–11 doi 10.1016/j.molcel.2009.11.016.

76. Shaffer AL, Wright G, Yang L, Powell J, Ngo V, Lamy L, et al. A library of gene expression signatures to illuminate normal and pathological lymphoid biology. Immunol Rev 2006;210:67–85 doi 10.1111/j.0105-2896.2006.00373.x.

77. Elemento O, Slonim N, Tavazoie S. A universal framework for regulatory element discovery across all genomes and data types. Mol Cell 2007;28(2):337–50 doi 10.1016/j.molcel.2007.09.027.

78. Schwammle V, Jensen ON. A simple and fast method to determine the parameters for fuzzy c-means cluster analysis. Bioinformatics 2010;26(22):2841–8 doi 10.1093/bioinformatics/btq534.

79. Corces MR, Trevino AE, Hamilton EG, Greenside PG, Sinnott-Armstrong NA, Vesuna S, et al. An improved ATAC-seq protocol reduces background and enables interrogation of frozen tissues. Nat Methods 2017;14(10):959–62 doi 10.1038/nmeth.4396.

80. Loven J, Hoke HA, Lin CY, Lau A, Orlando DA, Vakoc CR, et al. Selective inhibition of tumor oncogenes by disruption of super-enhancers. Cell 2013;153(2):320–34 doi 10.1016/j.cell.2013.03.036.

81. Meers MP, Bryson TD, Henikoff JG, Henikoff S. Improved CUT&RUN chromatin profiling tools. Elife 2019;8 doi 10.7554/eLife.46314.

82. van Galen P, Viny AD, Ram O, Ryan RJ, Cotton MJ, Donohue L, et al. A Multiplexed System for Quantitative Comparisons of Chromatin Landscapes. Mol Cell 2016;61(1):170–80 doi 10.1016/j.molcel.2015.11.003.

83. Ross-Innes CS, Stark R, Teschendorff AE, Holmes KA, Ali HR, Dunning MJ, et al. Differential oestrogen receptor binding is associated with clinical outcome in breast cancer. Nature 2012;481(7381):389–93 doi 10.1038/nature10730.

84. Liao Y, Smyth GK, Shi W. featureCounts: an efficient general purpose program for assigning sequence reads to genomic features. Bioinformatics 2014;30(7):923–30 doi 10.1093/bioinformatics/btt656.

85. Anders S, Huber W. Differential expression analysis for sequence count data. Genome Biol 2010;11(10):R106 doi 10.1186/gb-2010-11-10-r106.

86. Ramirez F, Ryan DP, Gruning B, Bhardwaj V, Kilpert F, Richter AS, et al. deepTools2: a next generation web server for deep-sequencing data analysis. Nucleic Acids Res 2016;44(W1):W160–5 doi 10.1093/nar/gkw257.

